# RATTLE: Reference-free reconstruction and quantification of transcriptomes from Nanopore sequencing

**DOI:** 10.1101/2020.02.08.939942

**Authors:** Ivan de la Rubia, Akanksha Srivastava, Wenjing Xue, Joel A Indi, Silvia Carbonell-Sala, Julien Lagarde, M Mar Albà, Eduardo Eyras

## Abstract

Nanopore sequencing enables the efficient and unbiased measurement of transcriptomes from any sample. However, current methods for transcript identification and quantification rely of mapping reads to a reference genome, which precludes the study of species with a partial or missing reference or the identification of disease-specific transcripts not readily identifiable from a reference. Here we present RATTLE, a tool to perform reference-free reconstruction and quantification of transcripts using only Nanopore reads. Using simulated data and experimental data from isoform spike-ins, human tissues, and cell lines, we show that RATTLE accurately determines transcript sequences and their abundances, and shows good scalability with the number of transcripts. RATTLE provides unprecedented access to transcriptomes from any sample and species without relying on a reference or additional technologies.

## Background

The direct interrogation of transcriptome has proven effective to study species without an available genome reference [1] or non-model organisms of ecological relevance [2], or to identify cancer-specific RNAs with diagnostic relevance [3]. However, these approaches have suffered from the limitations of the short sequencing reads, which lead to uncertainties in the reconstruction of transcripts [4]. Single-molecule long-read sequencing with Oxford Nanopore Technologies (ONT) enables the direct measurement of native RNA molecules, often identify complete known and novel transcripts [5], and accurately estimate transcript abundances [6]. These advantages coupled with the ease of sample preparation and the availability of portable platforms, makes this technology very compelling for the cost-effective and unbiased measurement of transcriptomes from any sample and species. However, current analysis methods rely on the comparison with a reference genome [5,7–9] or on the use of multiple sequencing technologies [10,11], which precludes the possibility of cost-effective studies. Methods for *de novo* DNA assembly [12–14] cannot be directly transferred to transcriptomes, since they do not model genes with multiple transcript isoforms. Similarly, methods developed for Illumina [1,15] rely on their high sequence accuracy, which is currently lacking in individual Nanopore reads. To address these challenges, we have developed RATTLE, a tool to perform reference-free reconstruction and quantification of transcripts using only Nanopore long reads. Using simulated data, isoform spike-ins, and sequencing data from human tissues and cell lines, we show that RATTLE is competitive at recovering transcript sequences and their abundances despite not using any information from the reference. RATTLE lays the foundation for a multitude of potential new applications of Nanopore transcriptomics.

## Results

### RATTLE algorithm

RATTLE starts by building read clusters that represent potential genes. To circumvent the quadratic complexity of an all-vs-all comparison of reads, RATTLE performs a greedy deterministic clustering using a two-step k-mer based similarity measure (Fig. 1). The first step consists of a fast comparison of k-mers (k=6) shared between every two reads (Supp. Fig. 1a), whereas the second step solves the Longest Increasing Subsequence (LIS) problem to find the longest list of collinear matching k-mers between a pair of reads, which defines the RATTLE similarity score (Supp. Fig. 1b). Clusters are greedily generated by comparing reads to a representative of each existing cluster at every step of the iteration. This process generates read clusters, where each cluster represents a potential gene containing reads originating from all possible transcript isoforms.

**Figure 1.**
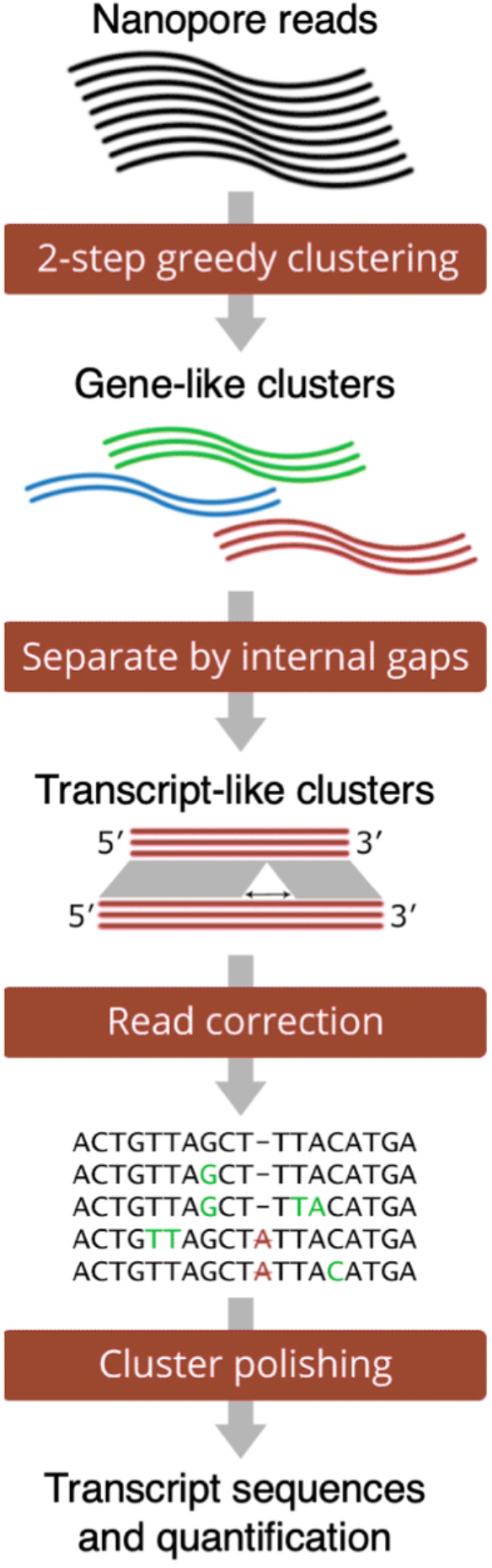
RATTLE proceeds through three main steps: clustering, error-correction, and transcript polishing. In the clustering step, gene-like and transcript-like clusters are produced. Error correction is performed at the level of transcript clusters. At the transcript- polishing step, transcript consensus sequences and their abundances are produced.

Gene clusters are subsequently split into sub-clusters defining candidate transcripts. These transcript clusters are built by determining whether every pair of reads in a gene cluster is more likely to originate from different transcript isoforms rather than from the same isoform. This is estimated using a threshold for the maximum variance allowed (--iso-max-variance) for the distribution of gap-lengths between the previously calculated co-linear matching k-mers (Supp. Fig. 1c). RATTLE performs error correction within each one of the transcript clusters by generating a block-guided multiple sequence alignment (MSA). Each read is assessed for error correction using the base quality for each base and the average quality of the consensus at each MSA column. RATTLE then builds the final transcripts after a polishing step to refine the cluster definitions. The final transcript sequence is obtained as a consensus from the final transcript cluster and the abundance is defined as the total read count in that cluster. More details are provided in the Methods section.

### Evaluation of read clustering into genes and transcripts

To illustrate the ability of RATTLE similarity score to perform read clustering, we simulated reads from multiple annotated transcripts using the read length distribution observed in a Nanopore cDNA sequencing run (Supp. Fig. 1d). RATTLE similarity score separated reads originating from different transcripts better than using the score based on a sequence alignment using Minimap2 [16], which is based on minimizers (Fig. 2a). This indicates that although minimizers provide an efficient comparison between sequences, RATTLE similarity score may be more robust for comparisons between error-prone reads. We further considered reads simulated from two transcripts from the same gene differing by only by an internal alternative exon (Supp. Fig. 1e). RATTLE similarity score separated reads originating from each of the two transcript isoforms better than using the number of matches in an alignment as similarity measure (Fig. 2b). Furthermore, we simulated reads from three pairs of transcripts differing by one internal alternative exon using 10 different exon lengths between 25 and 150, and for 7 different values of the -iso-max-variance parameter. The results showed that RATTLE correctly separated reads from two isoforms for alternative exons longer than 50-65 nt using --iso- max-variance 5 or higher (Supp. Fig. 2) (Supp. Table S1).

**Figure 2.**
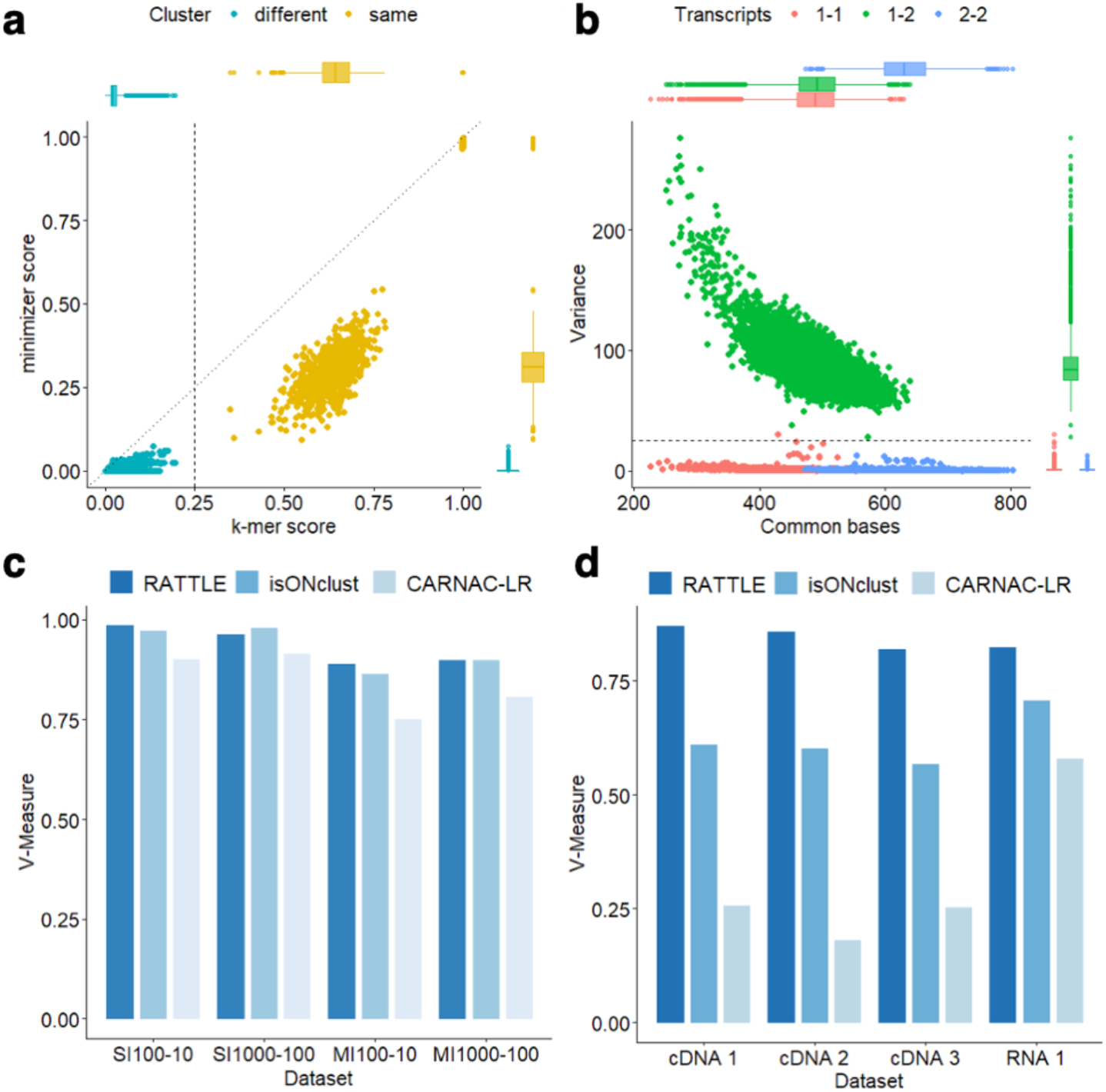
**(a)** Comparison of the RATTLE similarity score (x axis), based on the longest increasing subsequence, with a similarity score calculated from Minimap2 (y axis), using k=6. Each dot represents a comparison between a pair of reads simulated from the same (orange) or different (blue) transcripts from two different genes. The distribution of values for each comparison is given as box plots along each axis. **(b)** Number of common bases between two reads (x-axis) and variance in the distribution of gap-length differences between adjacent matching k-mers between the same two reads (y axis). Reads were simulated from two transcript isoforms from the same gene differing only by an internal exon (Supp. Fig. 1e). Each dot is colored according to whether the reads originated from the same transcript, 1-1 (red) or 2-2 (blue) or not, 1-2 (green). **(c)** Clustering accuracy of RATTLE, CARNAC, and isONclust in terms of the V-measure (y axis), using simulated reads. Simulations (x-axis) were performed with a single (SI) or with multiple (MI) transcript isoforms per gene, using a different number of total transcripts (*t*) and a different number of reads per transcript (*r*), indicated as SI*t*-*r* or MI*t*-*r*. Other accuracy metrics are provided in Supp. Table S2. **(d)** Clustering accuracy using spike-in isoform RNA variants (SIRVs) as reference. The plot shows the V-measure (y axis) for RATTLE, CARNAC, and isONclust using SIRV reads from four of the tested samples (x axis): three using the cDNA-seq protocol (cDNA1, 2, 3) and one using the dRNA protocol (RNA1) (Methods).

To test the accuracy in the identification of gene clusters, we compared RATTLE with two other methods that cluster long reads, CARNAC [16] and isONclust [17]. We built several reference datasets of simulated reads from multiple genes with one or more transcripts per gene and with a variable number of reads per transcript. RATTLE showed higher accuracy at recovering gene clusters in most of the comparisons and using different metrics (Fig. 2c) (Supp. Table S2). At the transcript level, RATTLE performed better than CARNAC and isONclust (Supp. Table S2). Moreover, RATTLE was faster than CARNAC and isONclust in all tested datasets (Supp. Table S2).

We further assessed the accuracy of RATTLE at recovering gene clusters using Spike-in isoform RNA Variants (SIRVs). The SIRV genome (SIRVome) is organized into 7 different genes containing a total set of 69 transcripts with known coordinates, sequences, and abundances. We performed 4 different experiments with added SIRVs: PCR-based cDNA sequencing (cDNA-seq) from human brain (two independent replicates, cDNA1, cDNA2) and human heart (cDNA3) samples, and direct RNA (dRNA-seq) from the same heart sample (RNA1). We first used the SIRV transcripts aggregated per gene to evaluate clustering at the gene level. RATTLE was faster and showed higher accuracy in the identification of SIRV genes compared with the other two methods (Fig. 2d) (Supp. Table S3). Similar results were found using dRNA and cDNA reads for SIRVs in mouse samples [6] (Supp. Table S3). CARNAC and isONclust appeared to be more sensitive to the dynamic range of SIRV abundances. Furthermore, RATTLE achieved high accuracy at clustering reads into SIRV transcripts using multiple metrics (Supp. Table S3).

### Evaluation of sequence accuracy after error correction

To test RATTLE accuracy in error correction in Nanopore reads without using a reference, we used the same SIRV reads and compared RATTLE results with methods developed for long-read self-correction. We used CONSENT [18] and isONcorrect [19], and the self-correction step from Canu [12], which has been used previously for the correction of cDNA reads [20]. Reads were mapped to the SIRVome with Minimap2 [21] before and after correction by each method. As a comparison, we included TranscriptClean [7] to correct the reads mapped to the genome reference without using the annotation. After correction, all methods improved the percentage identity with the SIRV transcript isoforms both for cDNA (Fig. 3a) and dRNA (Fig. 3b) reads (Supp. Fig. 3). All methods also showed a decrease in the error rates in cDNA (Fig. 3c) and dRNA (Fig. 3d) reads (Supp. Fig. 4). Both RATTLE and isONcorrect achieved results close to TranscriptClean, which had the best improvement over raw reads. TranscriptClean results were similar for cDNA and dRNA reads. However, self-correction with dRNA reads did not improve percentage identity and decrease error-rate as much as with cDNA reads. Notably, RATTLE was faster than all other methods in all tested samples, with runtimes ranging from 1.64 minutes (mins) (for 14,958 reads) to 123.9 mins (for 214,107 reads) (Supp. Table S4). Furthermore, RATTLE corrected approximately as many reads as TranscriptClean and isONcorrect, and more than CONSENT (Supp. Table S4). We also performed error correction using reads from the human transcriptome. RATTLE showed similar improvements over raw reads to isONcorrect and TranscriptClean (Figs. 3e and 3f) (Supp. Table S5).

**Figure 3.**
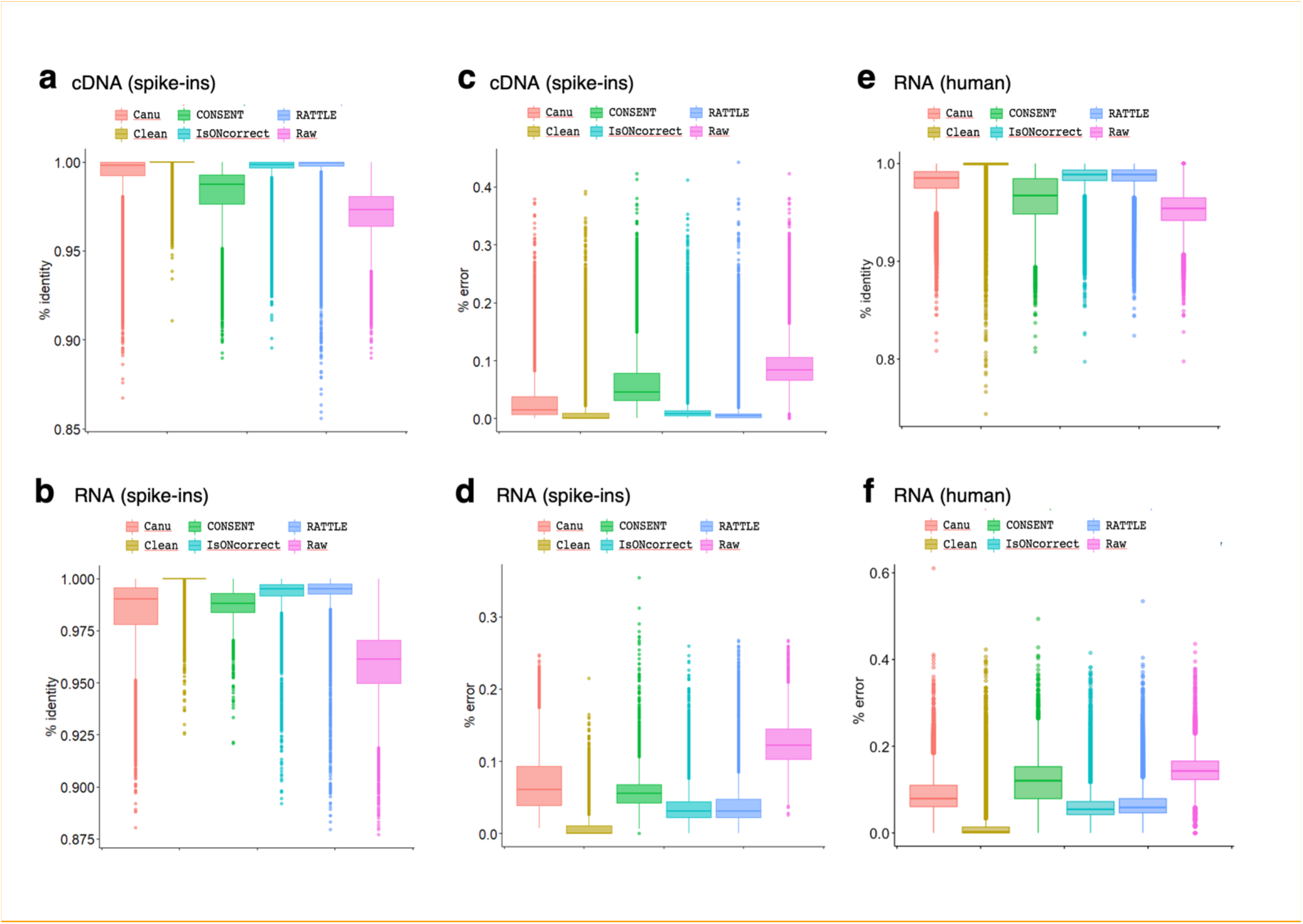
We show the percentage identity distributions of all SIRV reads before (Raw) and after correcting with RATTLE, CONSENT, isONcorrect, Canu, and TranscriptClean (Clean) for Nanopore cDNA (sample cDNA2) (a) and dRNA reads (sample RNA1) (b). Percentage identity was calculated as the number of nucleotide matches divided by the total length of the aligned region. Other samples are shown in Supp. Fig. 3. We also show the error rate distribution of SIRV reads before (Raw) and after correction with RATTLE, CONSENT, isONcorrect, Canu, and TranscriptClean (Clean) for the same cDNA (c) and RNA (d) samples. The error rate was calculated as the sum of insertions, deletions, and substitutions divided by the length of the read. Other samples are shown in Supp. Fig. 4. Finally, we show the distribution of percentage identity (e) and error rate (f) for reads from the human transcriptome before (Raw) and after correcting with RATTLE, CONSENT, isONcorrect, and TranscriptClean (Clean) for Human heart dRNA-seq (RNA1). Percentage identity and error rate was calculated as in (a-d).

**Figure 4.**
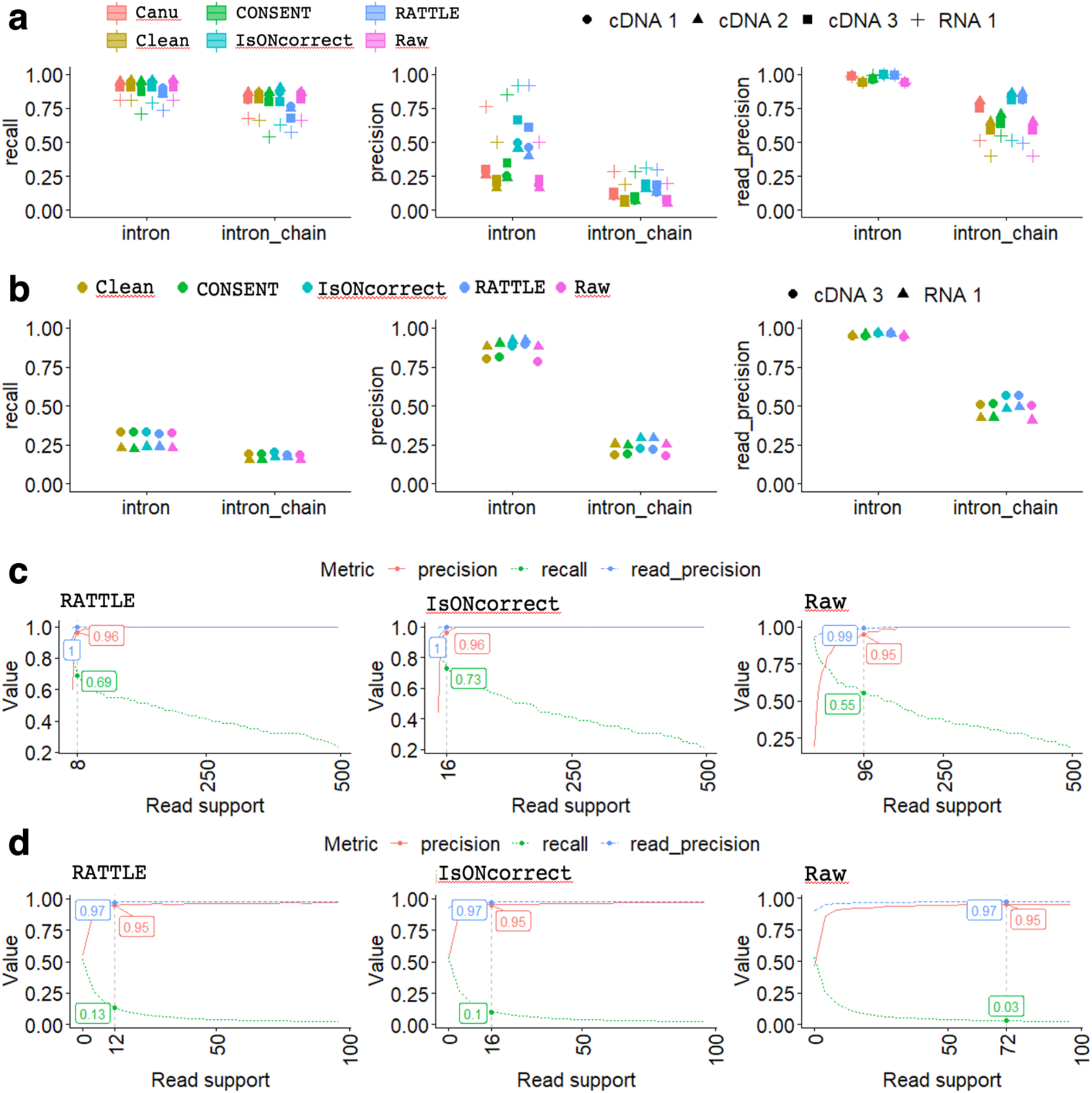
**(a)** Left panel: recall of unique SIRV introns and intron-chains obtained by mapping reads to the SIRV genome before (Raw) and after correction with RATTLE, CONSENT, Canu, isONcorrect, and TranscriptClean (Clean) for the dRNA-seq (RNA 1) and the cDNA (cDNA 1, 2, 3) samples. Recall was calculated as the fraction of unique annotated introns or intron-chains exactly found by each method. Middle panel: Precision values for introns and intron-chains for the same methods and datasets. Precision was calculated as the fraction of unique introns or intron-chains predicted by reads that matched exactly the SIRV annotation. Right panel: Read-precision for introns and intron-chains for the same methods and datasets. Read-precision was calculated as the fraction of all introns or intron-chains predicted in reads that corresponded to SIRV introns or intron chains. Similar plots for internal and external exons are given in Supp. Fig. 5. Only cases with >5 reads support were considered. **(b)** Same accuracy plots as in (a) for the same methods and datasets but using expressed Gencode transcripts as reference. **(c)** We plot the recall (green), precision (red) and read-precision (blue) of the SIRV introns (y axis), as a function of the number of minimum reads supporting the predictions (x axis). We indicate for each case the threshold at which a precision (red) of approximately 0.95 is achieved. For that threshold we indicate the corresponding recall (green) and read-precision (blue). The plot corresponds to the dataset cDNA2. Results for other samples are available in Supp. Fig. 7. **(d)** Same accuracy plots as in (c) but using expressed Gencode transcripts as reference. Results for other samples are available in Supp. Fig. 9.

### Evaluation of exon-intron structures after error correction

We next evaluated the ability to accurately recover exon-intron structures after mapping self-corrected reads to a genome reference. To this end, we used simple and easily interpretable metrics to determine the accuracy of the exon-intron structures. We first analyzed the exact matches to introns and intron-chains, defined as an ordered sequence of introns in an annotated transcript or mapped read (equivalent to FSM in the SQANTI nomenclature [22]). Using the SIRV reads and the SIRVome as reference, RATTLE recovered slightly fewer introns and intron-chains than other methods (Fig. 4a, left panel) but displayed precision values similar to isONcorrect and higher than the other methods (Fig. 4a, middle panel). The precision was generally low for all meethods, suggesting that despite the read correction, the proportion of unique false positive introns and intron-chains was still high for all methods. To further investigate this, we calculated a read-precision metric, defined as the proportion of reads supporting correctly identified annotated features. All tools showed an increase in read-precision with respect to precision (Fig. 4a, right panel), indicating that most of the wrongly predicted features are supported by a low number of reads. Indeed, false positive introns generally showed lower read support than true positives (Supp. Fig. 5).

We performed a similar analysis mapping sequencing reads to the human genome and testing the exact matches to the exon-intron structures of the human gene annotation. As no ground truth was available, we used as reference the exon-intron structures of Gencode transcripts that had evidence of expression in each sample. Compared to the analysis performed with the SIRVs, recall was relatively low for all methods, reflecting the higher transcriptome complexity of this dataset. RATTLE recovered a similar proportion of introns and intron-chains as the other tools, despite not using any information from the reference (Fig. 4b, left panel). In the case of precision (Fig. 4b, middle panel) and read-precision (Fig. 4b, right panel), RATTLE and isONcorrect achieved the highest values, suggesting that self-correction may help in discarding potential false positives.

We also performed comparisons to annotated exons, separated according to whether they were internal or external in the annotated transcript, using either SIRVs (Supp. Fig. 6) (Supp. Table S6) or the Gencode human transcriptome (Supp. Fig. 7) (Supp. Table S7). In general, the results recapitulated those observed for introns but we could also observe that external exons were less well recovered than internal ones, something expected given that the first 15 nucleotides at the 5’end of mRNAs are typically not recovered with ONT [5,23].

### Impact of the number of reads supporting a transcript

Our analyses indicated that it is fundamental to consider the number of supporting reads to establish the reliability of the predictions. To establish the accuracy in terms of read support, we calculated the recall, precision, and read-precision for the exact matches with SIRV introns at different thresholds of read support, and estimated the minimal read support needed to achieve a precision of 0.95 (i.e. 5% of the predictions would be false). All correction methods showed an improvement over the raw reads (Fig. 4c) (Supp. Fig. 8). RATTLE and isONcorrect were the methods that required the least read support to achieve 0.95 precision. Moreover, the recall at these thresholds for RATTLE and isONcorrect was also higher than for the other methods. We did the same analysis using the expressed Gencode human transcripts as reference and found again that RATTLE and isONcorrect required the least read support for 0.95 precision and achieved the highest recalls at these thresholds (Fig. 4d) (Supp. Fig. 9). These results indicate that RATTLE and isONcorrect generally require lower read support than other methods to ascertain the reliability of the introns defined by their read corrections.

### Evaluation of transcript abundance estimation

We next tested the accuracy of RATTLE transcripts to recover the known abundances of the SIRV transcript isoforms. We compared RATTLE predictions with StringTie2 [8], FLAIR [9], and TALON [23], which use the genome and annotation references to delineate transcript isoforms and their abundances. Despite not using any information from the SIRV genome or annotation, the correlation with SIRV isoform abundances achieved by RATTLE was comparable to the other methods (Fig. 5a) (Supp. Fig. 10) (Supp. Table S8). Interestingly, using a simple approach based on counting reads mapped to the transcripts achieved higher correlations than other methods for dRNA and cDNA reads (Supp. Fig. 10) (Supp. Table S8). In agreement with previous similar comparisons [6], the correlation for dRNA reads was generally higher than for cDNA for all methods (Supp. Fig. 10) (Supp. Table S8), indicating that dRNA reads produce better transcript abundance estimates than cDNA reads.

**Figure 5.**
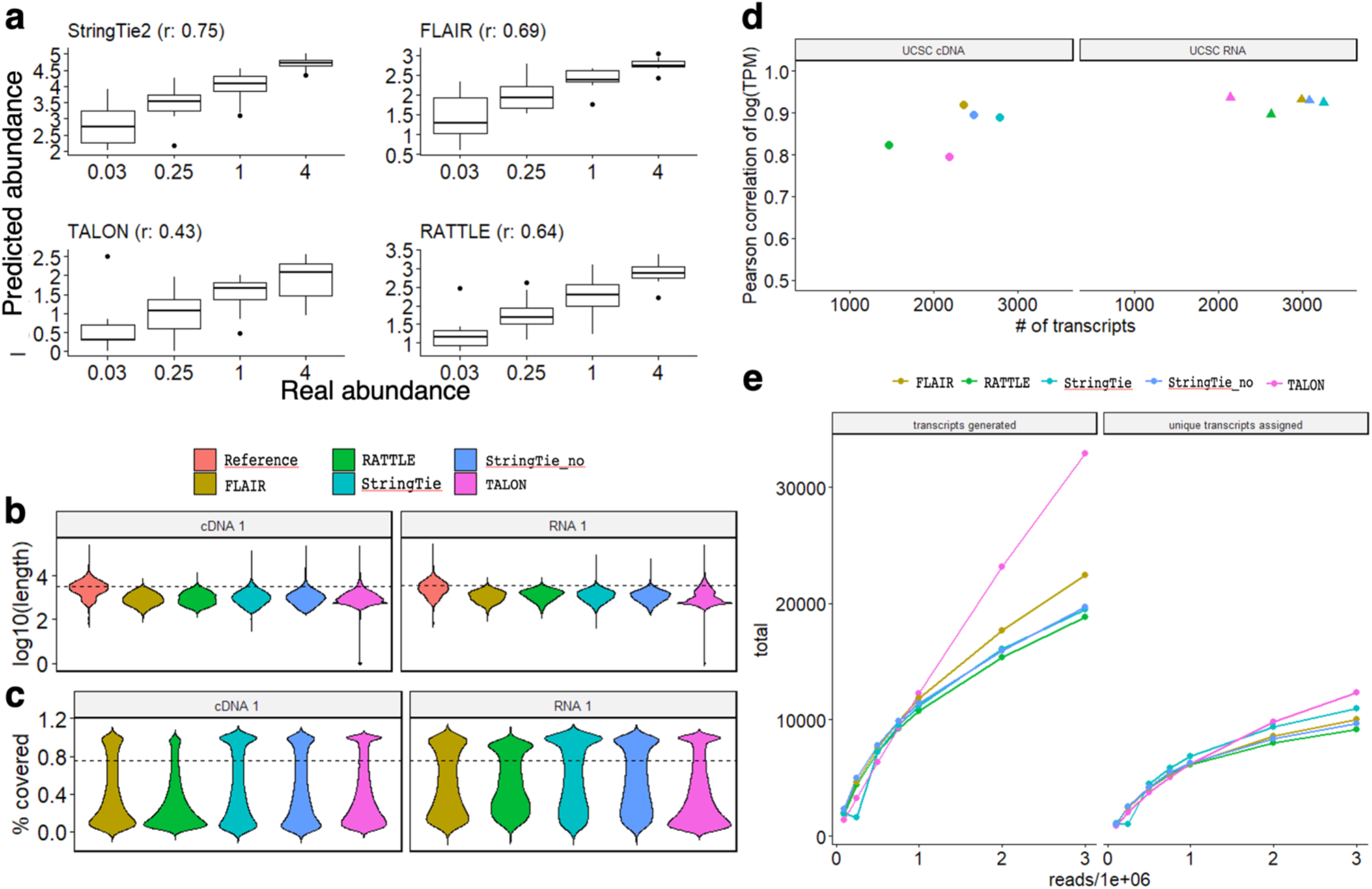
**(a)** Comparison of the transcript abundances (y axis) predicted by RATTLE, FLAIR, StringTie2, and TALON, with the abundances of the SIRV spike-in transcript isoforms (x axis). Each panel shows the Pearson correlation *r* for the comparison of the abundance values. Units on the y-axis vary according to the method: RATTLE provides abundances as read counts per million, like TALON and FLAIR. StringTie2 produces a TPM value. SIRV data corresponds to the RNA1 sample. Correlations for other datasets are provided in Supp. Fig. 10. **(b)** Distribution of the lengths (y-axis, log10 scale) of the transcripts predicted by each method compared with the transcripts from the reference (Gencode v29) using cDNA and dRNA. StringTie2 was run with (stringtie) and without (stringtie_no) the annotation. The dashed line indicates the median value for the reference transcripts. **(c)** Distribution of the proportion of the annotated transcript covered (y-axis) by the best matching predicted transcript from each method. The dashed line indicates 0.75. **(d)** Correlation (y-axis) of the predicted transcript abundances from two replicate cDNA and dRNA sequencing experiments from the Nanopore sequencing consortium. For each method, we give the number of predicted transcripts that were matched to the reference annotation in both replicates (x-axis). Colors are like for (b) and (c). **(e)** Saturation plot of transcripts using an incremental number of dRNA reads as input. Left panel: for each input (x-axis), we give the total number of transcripts predicted by each method (y-axis) with an expression of >5 reads. Right panel: for each input (x-axis), we give the total number of annotated transcripts matched by the predicted transcripts from the left panel with a coverage of at least 75%. The number of transcripts obtained by each method and input is given in Supp. Table S10.

We next decided to study the properties of the RATTLE outputs using human transcriptomes. As no ground truth was available for the human transcriptome, we had to devise a way to perform controlled comparisons. Using the same human brain and heart samples as before, we first studied the length distributions of the transcripts predicted by each method. RATTLE transcripts showed similar length distributions to reference-based methods (Fig. 5b) (Supp. Fig. 11). Moreover, all methods produced transcripts that were generally shorter than the annotation (Gencode v29) (median length indicated as a dashed line in Fig. 5b). To compare all tools on similar conditions, we remapped the transcripts predicted by each tool directly to the Gencode transcriptome using Minimap2. Looking at the coverage of the annotated transcripts by the matching predicted transcripts, all tools showed a bimodal distribution (Fig. 5c) (Supp. Fig. 12). This behaviour was similar across the tools for dRNA, but RATTLE and FLAIR showed lower coverage values for cDNA. Our analyses also suggested that transcript matches with a coverage of at least 75% (dashed line in Fig. 5c) may provide a good correspondence between predictions and annotations. We thus used this strategy to determine the reproducibility of the abundances predicted by each method across replicates of cDNA and dRNA sequencing from a lymphoblastoid cell line [5]. The transcripts predicted by RATTLE, FLAIR, StringTie2, and TALON were directly mapped to the human transcriptome annotation, keeping only one-to-one matches with at least 75% coverage (Methods). The abundance of the predicted transcripts assigned to the same annotated transcript was then compared across replicates. Although RATTLE generally recovered fewer transcripts (Supp. Table S9), it showed similar correlation values to the reference-based methods (Fig. 5d). These correlation values were also stable across other coverage cutoffs (Supp. Fig. 13). Moreover, the correlation values for all methods were generally higher for dRNA than for cDNA reads (Fig. 3d) (Supp. Table S9).

### RATTLE scalability

To investigate the relationship between the number of predicted transcripts and the number of input reads, we used data from multiple dRNA runs in the same cell line to generate input datasets of size 100k, 250k, 500k, 750k, 1M, 2M, and 3M reads. The total number of transcripts increased with the number of reads although in a sub-linear manner, except for Talon (Fig. 5e left panel) (Supp. Table S10). We further measured the number of predicted transcripts that had a matched annotated transcript using the same approach as above. The curves relating the number of transcripts and the number of input reads showed a tendency to saturate and were similar for the different methods (Fig. 5e right panel). We further compared RATTLE and RNA-Bloom [24] using input datasets from 100k to 3M reads from a different experiment (Methods). This analysis showed that RATTLE transcripts are longer than RNA-Bloom transcripts for all input sizes and converged to the expected length of the annotation (Supp. Fig. 14). To further compare both tools, we calculated the number of transcripts produced by both methods for each input size. The number of RATTLE transcripts increased in a sublinear manner, whereas the number of RNA-Bloom transcripts increased at a much faster rate with the input reads (Supp. Fig. 15)

Finally, we studied the running time and memory usage for all RATTLE steps, i.e., clustering, read correction, and transcript polishing. We first tested five different configurations for the parameters of the clustering step with 3 different input sizes, 100k, 250k, 500k reads, using the SIRV transcriptome to test the accuracy of each run (Supp. Table S11). Using different metrics, these tests showed that RATTLE recovered SIRV genes and transcripts with similar high accuracy across all parameter configurations and input sizes. We also observed that the memory usage was only dependent on the number of input reads. On the other hand, the running time was slightly lower for parameter configurations that resulted in fewer iterations of the greedy clustering algorithm (see configuration c5 in Supp. Table S11).

Next, we calculated the memory usage and running times for input datasets of 100k, 250k, 500k, 750k, 1M, 2M, and 3M reads. The read correction step showed a larger memory peak than the clustering step for an input size of <1M reads (Fig. 6a), similarly to the pattern previously observed with the SIRV reads. However, the clustering step was close to a linear form for the clustering step and showed higher memory usage beyond 1M input reads. The running time for all RATTLE steps together was slightly over linear, and much faster than quadratic time (Fig. 6b) (Supp. Table S12). Although RNA-Bloom was generally faster for these inputs, it required much higher memory (Supp. Table S12).

**Figure 6.**
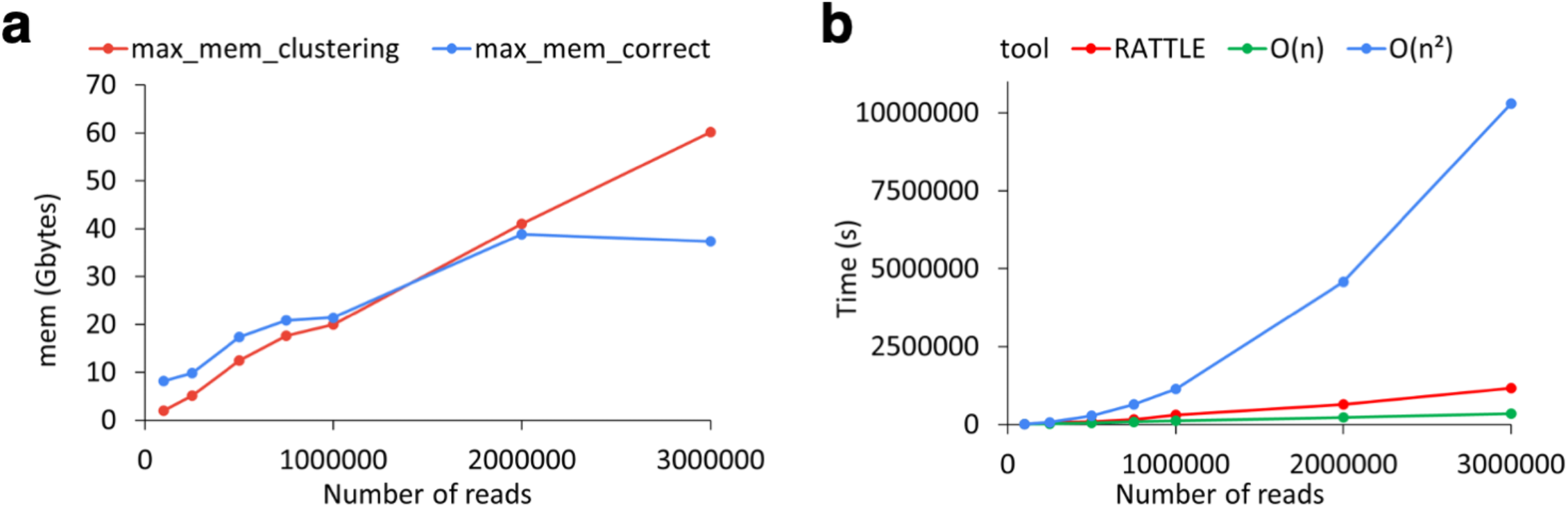
**(a)** Maximum memory usage in Gigabytes (y axis) as a function of the number of input reads (x axis) for RATTLE clustering (red) and read correction (blue) steps. **(b)** Sum of the CPU time in seconds (y axis) for the RATTLE clustering, correction, and polishing steps for an increasing number of input reads (x axis). RATTLE time (red) is compared with the linear (green) and quadratic (blue) times. The O(n) and O(n^2^) graphs are extrapolated from the first value. For (a) and (b), RATTLE was run with the parameter configuration c5 described in (Supp. Table S11). Memory and timing values are given in Supp. Table S12.

## Discussion

Our analyses provide evidence that despite the current limitations of using ONT reads without a reference genome, RATTLE represents a significant breakthrough in the capacity to perform reference-free transcriptomics with Nanopore reads, without using additional technologies. Using multiple data types and tests, RATTLE shows a competitive accuracy at building genes and transcripts, as well as at estimating their abundances. RATTLE generally achieved a high precision in the exon-intron structures produced by corrected reads and predicted transcripts, which will be fundamental to reliably identify new genes and transcript isoforms in samples without an available reference. In our tests, RATTLE performed comparably to reference-based methods, and self-correction did not impact the ability of RATTLE to recover exon-intron structures with high precision.

Several limitations remain for reference-free long-read transcriptomics. RATTLE generally recovered fewer known transcripts compared to the reference-based methods. This indicates a limitation in the resolution of clusters when real biological differences are comparable to sequencing errors in reads. In particular, we observed that there is a lower limit in the size of alternative exons that RATTLE can separate between pairs of individual reads, which impacts the identification of transcript isoforms and their abundances. We also observed that quantification with cDNA reads, in comparison with dRNA reads, showed in general high variability across methods, lower accuracy in comparison with the true abundances, and lower correlation across replicates. Our analyses also indicated that simply counting mapped reads can accomplish high accuracy. This suggests that, despite having lower sequence accuracy, dRNA may capture better the lengths and abundances of RNA molecules, and hence may be more suitable for reference-free transcriptomics. Given that abundance is a relative measure and as the length and accuracy of dRNA reads improve with time, accurate quantification might be achieved by simply counting reads mapped to transcripts. In turn, this will also improve the reference-free estimation of abundances with RATTLE.

In comparison with other read clustering and error correction methods, RATTLE was generally faster. RATTLE algorithms and data structures facilitate the efficient comparison of reads for clustering, and the storage and processing for transcript prediction. Although other approaches may be faster at clustering reads into candidate transcripts, RATTLE achieves higher accuracy in terms of transcript lengths and abundances. RATTLE thus provides competitive reference-free transcriptome analysis for the most common outputs from a MinION platform (500k-3M reads). Further improvements may be required to make a reference-free approach competitive for much larger input datasets, such as those from the PromethION platform, which may yield over 20M dRNA reads.

RATTLE modularity, with the ability to parameterize each step, means it can be easily adapted to any sample type. Additionally, RATTLE rich output, including information about the predicted transcripts and genes, as well as the reads used to build each transcript, will prove valuable in downstream analyses, including the study of differential transcript usage [25], the analysis of single-cell long-read sequencing [26], and the identification of RNA modifications in non-model species [27]. RATTLE lays the foundation of exciting developments in long-read transcriptomics. Considering the rapid growth of the interest in ONT RNA sequencing in a variety of new samples and systems, RATTLE can become a valuable tool for the scientific community.

## Data availability

Nanopore sequencing data generated in this study has been deposited in the European Nucleotide Archive (ENA) under the umbrella study PRJEB40410 (https://www.ebi.ac.uk/ena/browser/view/PRJEB40410) including the FASTQ files (https://www.ebi.ac.uk/ena/browser/view/PRJEB39835) and signal data (https://www.ebi.ac.uk/ena/browser/view/PRJEB40335). Datasets from mouse used in this study were obtained ENA under study accession PRJEB27590 (http://www.ebi.ac.uk/ena/data/view/PRJEB27590) (runs ERR2680375 and ERR2680377), and from the Nanopore sequencing consortium from https://github.com/nanopore-wgs-consortium/NA12878 (under nanopore-human-transcriptome). Direct RNA sequencing data from HEK293 cells was obtained from ENA, under the accession PRJEB40872 (http://www.ebi.ac.uk/ena/data/view/PRJEB40872). Data files with the simulated sets of reads, as well as the output files generated by the different assembly programs have been deposited in figshare (https://figshare.com/projects/RATTLE_Paper_Data/113580).

## Software availability

RATTLE is written in C++ and is available at https://github.com/comprna/RATTLE under the GPL-3.0 license.

## Methods

### RATTLE clustering algorithm

Before running RATTLE, reads were pre-processed with porechop (https://github.com/rrwick/Porechop) and those of length 150nt or shorter were filtered out. RATTLE starts by performing a greedy clustering. RATTLE sorts the reads in descending order by their length and processes them one at a time in that order. In the first iteration, RATTLE creates a new cluster with the first unclustered read. All subsequent unclustered reads are then compared against each existing cluster and assigned greedily if the scores are above a certain threshold. Otherwise, a new cluster is created. In each subsequent iteration, the threshold is decreased by a step change, and clusters are created or merged as initially. This process is continued until the threshold used is reduced to a limit. To ensure fast computation and circumvent the quadratic time complexity of an all-vs-all comparison, comparisons are performed using a representative read from each cluster, which is defined by the position in the ranking of read lengths within the cluster and can be set as a parameter by the user. In our analyses, we used the read at the position 0.15 x (number of reads in the cluster). At each iteration, and for each existing cluster, all other clusters and unclustered reads are compared to this cluster using the cluster representative. When two clusters are similar above the set thresholds, a new cluster is formed with the reads from these two clusters.

To reduce memory usage and for efficient calculation, reads are compared using a two-step similarity calculation. The first one provides a fast approximate comparison using bit-vectors, whereas the second one provides a more sensitive comparison if a threshold is passed. For the bit-vector comparison, sequence k-mers in reads are hashed into 32-bit integers with the hashing function H(A)=0, H(C)=1, H(G)=2, H(T)=3, such that for any k-mer s=b1…bk, the hash H(s) = 4^k-1^H(b1) + 4^k-2^H(b2) + … + H(bk) maps each k-mer to a unique position of a vector that is set to 1, i.e. a k-mer bit-vector. Each read is converted in this way into a bit-vector where the positions in the vector corresponding to the hashed k-mer in the read are set to 1. The first similarity score between a pair of reads is obtained through an AND operation on their bit-vectors and calculated as the number of common k-mers divided by the maximum number of k-mers in either read. Extraction and hashing of k-mers is performed only once per read in linear time, and the vector operations are performed in constant time.

If this first score is above a previously set threshold, a second similarity calculation is performed. For the second metric, all k-mers from both reads are extracted. Now k can be chosen on the command line, and k- mers are hashed as before. The intersection of the k-mers from both reads and their positions in each read are used to generate a list of triplets (*s, p1, p2*), where *s* is the hashed k-mer, *p1* is the position of this k-mer in the first read, and *p2* is the position of the same k-mer in the second read. These triplets are then sorted by *p1* and the Longest Increasing Subsequence (LIS) problem is solved with dynamic programming for *p2*. This produces the longest set of common co-linear k-mers between a pair of reads. The similarity value is defined as the number of bases covered by these co-linear common k-mers over the length of the shortest read in the pair. If the orientation for cDNA reads is unknown [28], RATTLE tests both relative orientations for each pair of reads by default, or only the given strand if the --rna option is used. As a result, all reads within a cluster have the same orientation.

The number of iterations to be performed for clustering is specified in the command line by setting the initial (defined by -B) and final (defined by -b) thresholds for the first bitvector-based score (default 0.4 to 0.2) and a decreasing step change (defined by -f) (default 0.05). For default parameters, iterations are thus performed for thresholds 0.4, 0.35, 0.3, 0.25, and 0.2. A final comparison is done using a threshold of 0.0, i.e., all remaining singletons and all cluster representatives are compared to each other using the LIS-based score. The LIS-based score threshold remains fixed over the entire clustering process. In our analyses, it was required to be 0.2 or larger. Analyses shown here were carried out for k=10 (can be changed with -k). Please see https://github.com/comprna/RATTLE for a complete description of all parameters.

### RATTLE transcript-cluster identification and error correction

Initial read clusters produced by the algorithm described above are considered to approximately correspond to genes, i.e., gene-clusters. Reads within each cluster are then separated into subclusters according to whether they are likely to originate from different transcript isoforms to form transcript-clusters. RATTLE considers the relative distances between co-linear k-mers calculated from the LIS-based score. Two reads in the same gene-cluster are separated into different transcript-clusters if the distribution of the relative distances between co-linear matching k-mers has a variance greater than a given threshold. That is, if co-linear matching k-mers calculated from the LIS algorithm show relative distances that would be compatible with a difference in exon content. The value 25 was used for the analyses, but it can be also modified as an input parameter.

RATTLE performs read correction within each transcript-cluster in two steps. First, each cluster with N reads is separated into blocks, each with R reads. In our analyses, R was set to 200. If R≤N<2R, the cluster is split in half, and if N<R, we take a single block. To avoid length bias, blocks are built in parallel from the reads in the cluster sorted by length: to build K blocks, block 1 is made from reads in positions 1, K+1, 2K+1, …; block 2 from reads in positions 2, K+2, 2K+2, … ; and block K from reads in positions K, 2K, 3K, …. A multiple sequence alignment (MSA) is then built from each block using SIMD partial order alignment (SPOA) (https://github.com/rvaser/spoa) [29]. A consensus from each column in the MSA is then extracted in the following way: for each read and each base of the read, the base is changed to the consensus if the consensus base occurs with at least 60% frequency, but not if this base has an error probability (obtained from the FASTQ file) less than or equal to 1/3 times the average for the consensus base in that position of the alignment. Indels are treated similarly, but without the error constraint. This is only performed using aligned positions. That is, for each base in a given read, other reads of the MSA contribute to the correction if they have a base or an internal gap in that position. The consensus sequences from all blocks are then realigned with SPOA to obtain a final MSA for the transcript-cluster and an associated consensus is obtained as before. Only transcript-clusters with a minimum number of reads are corrected and considered for further analysis. Here we used transcript-clusters with more than 5 reads. The frequency of the consensus, error-probability cutoff, and the minimum number of reads for a transcript cluster can be set up as input parameters. We observed that in MSAs many reads had a few bases wrongly aligned at the terminal regions. To fix these cases so that they do not affect the correction step, RATTLE identifies small blocks (<10nt) followed by large gaps (larger than or equal to 20 positions) at both ends of each aligned read and removes them. A block is defined as a subsequence that has gaps shorter than or equal to 3nt. RATTLE keeps removing blocks that satisfy these conditions at both ends of every aligned read until there no more such blocks are found.

### RATTLE transcript polishing and quantification

To define the final list of transcripts, RATTLE performs a final polishing step of the transcript-cluster definitions. This is done to correct possible splitting of reads originating from the same transcript into different transcript-clusters (under-clustering). For this, RATTLE uses the same 2-step greedy clustering described above on the transcript-clusters using the representative read from each cluster in the comparison. In each of the resulting clusters, an MSA column consensus is calculated, with abundance given by all the reads contained in the final cluster. Additionally, the transcripts are given a gene ID that corresponds to the gene-clusters to which they belong. When two transcript clusters are merged, if they are part of the same original gene-cluster, the resulting transcript stays in the same gene. If they are part of different genes, the gene with more transcripts absorbs the transcripts from the other gene to become one single gene.

RATTLE outputs different files at different stages of its execution. In the clustering step, it can either output gene-clusters or transcript-clusters in binary files. These files can then be used to extract a CSV file containing each read ID and the cluster it belongs to. The same binary files are also used for the correction step, which outputs three files: one with the corrected reads, one with those that are left uncorrected, and one containing the consensus sequence for each cluster from the input (in FASTQ format). Finally, the transcript-cluster polishing step receives as input the consensus sequences from the correction step and outputs a new file in FASTQ format with the final transcriptome. The quantification of each transcript is included in the header line. The quality per base in each FASTQ entry is calculated as the average of the PHRED quality scores from the bases in each column of the MSA in the transcript-cluster.

### Simulated reads and clusters

We developed a wrapper script (available at https://github.com/comprna/RATTLE) for DeepSimulator [30] to simulate a specific number of reads per transcript considering a read length distribution. The length distribution was calculated from a human cDNA sequencing sample from the Nanopore consortium [5]. To simulate the read sequences, we used the Gencode transcript annotation (v29), filtering out pseudogenes and genes from non-standard chromosomes, and removing transcripts that showed > 95% percentage identity with other transcripts using CD-HIT [31]. We then randomly selected different numbers of genes and transcripts to simulate reads. We considered genes with one single transcript isoform (SI), or genes with multiple isoforms (MI). For each case, various datasets were simulated using different numbers of reads per transcript and different numbers of transcripts per gene. To determine the accuracy of the clustering we used the adjusted rand index, which is a measure of the similarity between two cluster sets corrected for chance [32]. Additionally, we used homogeneity, completeness, and the V-measure [33]. Homogeneity is maximal when each cluster contains only elements of the same class. Completeness is maximal when all the elements of a single class are in the same cluster. The V-measure is the harmonic mean of completeness and homogeneity. We compared the clusters predicted by each method with the simulated clusters as the reference set. We run isONclust [17] (options: --ont --t 24), CARNAC-LR [16] with Minimap2 overlaps (options: -t 24 -x ava-ont), and RATTLE clustering (options: -t 24 -k 10 -s 0.20 –v 1000000 –iso-score-threshold 0.30 –iso-kmer-size 11 –iso-max-variance 25 –p 0.15).

### MinION sequencing with SIRVs

We performed Nanopore sequencing on two total RNA samples from human brain (Ambion - product num. AM7962; lot num. 1887911) and heart (Ambion - product num. AM7966; lot num. 1866106). Unless otherwise noted, kit-based protocols described below followed the manufacturer’s instructions. Regular quality controls using qBIT, nanodrop and Bioanalyzer were performed according to the manufacturer’s protocols to assess the length and the concentration of the samples. rRNA depletion was performed using Ribo-Zero rRNA Removal Kit Human/Mouse/Rat (Epicentre - Illumina). 12 ug of total RNA from each sample were prepared and divided into 3 aliquots (4 ug of total RNA each). 8ul of a 1:100 dilution (1 ng total) of synthetic controls (E2 mix lot number 001418 from SIRV-set, Lexogen) were added to each total RNA aliquot. The resulting ribosomal depleted RNAs were purified using 1.8X Agencourt RNAClean XP beads (Beckman Coulter). Samples were finally resuspended with 11 ul of RNA-free water and stored at -80°C. The cDNA was prepared using 50 ng of rRNA depleted RNA. The Takara Bio cDNA synthesis kit based on SMART (Switching Mechanism at 5’ End of RNA Template) technology coupled with PCR amplification was used to generate high yields of full-length double-stranded cDNA.

The sequencing libraries were prepared using 1 ug of full-length double-stranded cDNA following the standard ONT protocol SQK-LSK108 for an aliquot of the brain sample (cDNA 2), protocol SQK-LSK109 for 1 aliquot of the brain sample (cDNA 3) and 1 aliquot of the heart sample (cDNA 4). The direct RNA sequencing library was prepared using 500 ng of previously prepared ribosomal depleted sample (RiboZero kit, catalog num. MRZH11124, Epicentre-Illumina) from heart (RNA 2) with standard direct RNA ONT protocol SQK-RNA002, following manufacturer’s instructions, including the RT step. The final libraries were loaded on an R9.4.1 flowcell, and standard ONT scripts available in MinKNOW were used for a total of 48 hours run for each flowcell. ONT sequencing data was basecalled using Guppy 2.3.1+9514fbc (options: -- qscore_filtering --min_qscore 8 --flowcell FLO-MIN106 --kit <kit> --records_per_fastq 0 --recursive -- cpu_threads_per_caller 14 --num_callers 1), where <kit> is the corresponding protocol, as specified above (SQK-LSK109, SQK-LSK108 or SQK-RNA002).

To select reads corresponding to SIRVs, we run porechop (https://github.com/rrwick/Porechop) and mapped the reads to the SIRV genome (SIRVome) with Minimap2 (options: -t 24 -cx splice --splice-flank=no -- secondary=no). To extract the subset of reads with a hit on the SIRVome and being at least 150nt in length we used seqtk (https://github.com/lh3/seqtk, option subseq). We also used data from cDNA (ERR2680377) (cDNA_br_mm) and direct RNA (ERR2680375) (RNA_br_mm) sequencing of mouse brain including the E2 SIRVs [6], as well as samples from the Nanopore sequencing consortium [5].

### Clustering accuracy analysis

We first built SIRV isoform clusters by mapping reads to SIRV isoforms with Minimap2 and selecting for each read the SIRV isoform with the best mapping score. All reads that mapped to the same SIRV gene were then considered a gene-cluster. We then clustered reads with RATTLE, CARNAC, and isONclust and measured the accuracy of the predicted clusters by comparing them with the built SIRV gene clusters using the same metrics as with the simulated data. We used subsets of 25,000 reads (subsampled with seqtk) from each sample since we could not make CARNAC run for some of the complete datasets.

### Assessment of error correction accuracy

Reads were mapped to the SIRV transcripts with Minimap2 before and after read correction. Here we used the complete dataset of SIRV reads. Each read was assigned to the best matching transcript according to the mapping score. From the CIGAR string of the SAM output, the error rate was calculated as the sum of insertions, deletions, and substitutions divided by the length of the read, and the percentage identity as the number of nucleotide matches divided by the total length of the aligned region. We compared RATTLE (options: -t24 –g 0.3 –m 0.3 –s 200) with CONSENT [18] (options: consent-correct --type ONT), Canu [12] (options: minReadLength=200 stopOnLowCoverage=0.10 executiveMemory=16 executiveThreads=24), isONcorrect [19] (options: --t 24), and TranscriptClean [7]. TranscriptClean was run with default parameters using as input the reads mapped with Minimap2 (options: -t 12 -cx splice --splice-flank=no --secondary=no), but with no exon-intron information from the SIRV annotation.

To perform the accuracy analysis of the SIRV annotation features, PAF files from the mapping were compared with the annotation using ssCheck (available at https://github.com/comprna/RATTLE/). ssCheck works by comparing annotation features in the mapped reads with the annotation and calculates the number of unique features as well as the total number of features predicted in the mapped reads. As annotation features, we used introns, intron-chains. internal exons and external exons. An intron-chain was defined as an ordered sequence of introns in an annotated transcript or mapped read. The recall was calculated as the fraction of unique annotated features correctly found; precision was calculated as the fraction of unique predicted features that were in the annotation and read-precision was calculated as the fraction of the total number of predicted features in reads that corresponded to annotated features. Read-precision is affected by abundance levels but better reflects the accuracy per read. We developed ssCheck to be able to calculate the read-precision, since gffcompare (https://ccb.jhu.edu/software/stringtie/gffcompare.shtml) merges identical intron chains with different identifiers, which precludes this calculation. Additionally, ssCheck calculates accuracy metrics for internal and external exons, which was not possible with gffcompare.

To perform the comparison with the human transcriptome we considered the Gencode transcripts (v29) after removing pseudogenes, genes from non-standard chromosomes, and transcripts with >95% percentage identity with other transcripts. We only used transcripts that had evidence of expression of >5 reads from mapping the reads to the transcriptome using Minimap2 (options: -t 24 -x map-ont --secondary=no) to avoid spurious matches to transcripts with shared sequence but without evidence of expression in the sample, as done previously [34]. We then mapped the reads corrected by each method to the human genome with Minimap2 (options: -t 24 -cx splice --secondary=no). Considering the various genomic features in the expressed annotated transcriptome (introns, intron-chains, internal exons, external exons), we calculated the recall, precision, and read-precision with ssCheck as before.

### Assessment of transcriptome sequence and abundance accuracy

We used FLAIR [9] (options: align, correct, collapse, quantify –tpm), StringTie2 [8] (options: -p 24 –L) and TALON [23] (talon_initialize_database, talon, talon_summarize, talon_abundance, talon_create_GTF) with the cDNA and dRNA reads mapped to the SIRV genome with Minimap2 (options: -t 24 –ax splice –splice-flank=no –secondary=no, with –MD tag for TALON). These methods perform read correction (FLAIR and TALON) and transcript quantification (FLAIR, StringTie2 and TALON) of annotated and novel transcripts using the mapped reads with the help of the annotation. For the same samples, RATTLE was run for clustering (options: -t 24, -k 10, -s 0.20 –v 1000000 –iso-score-threshold 0.30 –iso-kmer-size 11 –iso-max-variance 25 –p 0.15), read correction (options: -t24 –g 0.3 –m 0.3 –s 200) and transcript polishing (options: -t 24). As an additional comparison, we mapped raw reads directly to SIRV isoforms with Minimap2 (options: -ax map-ont -t24) and estimated abundances with NanoCount [35], which assigns reads to isoforms with an expectation-maximization (EM) algorithm. We also assigned reads directly to SIRV isoforms by mapping them with Minimap2 (options: -t 24 -cx map-ont --secondary=no) and simply counting the number of mapped reads per transcript. FLAIR, StringTie2, and TALON provide the SIRV isoform ID with the predicted abundance. If more than one predicted transcript mapped to the same annotation, we only considered the prediction with the highest abundance. To assess the accuracy of RATTLE, we matched transcripts predicted by RATTLE to the SIRV isoforms using Minimap2 (options: -cx map-ont --secondary=no), assigning each SIRV to the best matching RATTLE transcript. If more than one transcript matched the same SIRV isoform, we selected the RATTLE transcript with the highest abundance.

To assess the accuracy of the transcriptome in terms of exon-intron structures, we compared the predicted transcripts mapped to the genome with the Gencode transcripts with evidence of expression (>5 reads) in the same sample. To make this analysis comparable across methods, all predicted transcriptomes were mapped against the genome with Minimap2 with Minimap2 (options: -t 24 -cx splice --secondary=no). We then calculated the exact matches of the predicted intron-exon structures (introns, intron chains, internal exons, external exons) with the annotation for that sample using ssCheck as before.

### Correlation of human transcriptomes

We used data from cDNA and dRNA sequencing from the Nanopore consortium [5]: the samples from Johns Hopkins (cDNA 2 replicates, dRNA 2 replicates) and UCSC (cDNA 2 replicates, dRNA 2 replicates). The resulting transcriptome from each method, FLAIR, TALON, RATTLE, and StringTie2 (with and without using the annotation), was then mapped to the expressed annotated transcripts using Minimap2 (options: -t 24 -x map-ont --secondary=no). To perform an accurate assignment of predicted to annotated transcripts, for each predicted transcript we only kept the best match if it covered at least 75% of the annotated transcript. Moreover, for each annotated transcript, we only assigned the best matching predicted transcript. If more than one passed the filter, we assigned the predicted transcript with the highest abundance. That is, the assignments of predicted and annotated transcripts were one-to-one. The correlation was calculated using the abundances of the predicted transcripts in the two replicates matched to the same annotated transcript.

### Scalability analysis

We used 30 pooled direct RNA sequencing runs from the Nanopore sequencing consortium [5] (https://github.com/nanopore-wgs-consortium/NA12878/blob/master/RNA.md) and generated subsamples of 100k, 250k, 500k, 750k, 1M, 2M, and 3M reads, using with seqtk (https://github.com/lh3/seqtk). We run FLAIR, TALON and StringTie (with and without annotation) as before on all these samples using the same options as before. RATTLE was run in all these samples (options: -B 0.5 -b 0.3 -f 0.2 --iso) using a machine with 24 cores. We first considered the total number of transcripts predicted by each method for each input size with an expression >5 reads. For each input size we also performed a one-to-one assignment of the predicted transcripts to the transcriptome annotation as described above to calculate the total number of identified annotated transcripts for each input size. We also run RATTLE (options: -B 0.5 -b 0.3 -f 0.2 --iso) and RNA-Bloom [24] (option: -long) in datasets of size 100k, 250k, 500k, 750k, 1M, 2M, and 3M reads, by subsampling reads from three dRNA replicates from HEK293 cells [36]. The memory usage and running times for RATTLE (Supp. Table S12) were obtained for these input datasets.

## Acknowledgments

We thank Roderic Guigó for providing access to the samples with SIRVs. We also thank the Nanopore sequencing consortium for providing access to their Nanopore transcriptomic sequencing runs. We acknowledge funding support funding from the Australian Research Council with ID DP210102385; from the Spanish Government and FEDER with grants BFU2015-65235-P (MMA), BIO2017-85364-R (EE), PGC2018-094091-B-I00 (MMA); and from the Catalan Government (AGAUR) with grant SGR2017-1020 (MMA). IdlR was supported by grant PRE2018-083413 from the Spanish Government.

## Author contributions

EE and IdlR designed the algorithms in RATTLE with inputs from MMA. IdlR prototyped and implemented the algorithms. IdlR carried out the benchmarking analyses with some contributions from JAI. AS and WX contributed to the software development. SCS and JL generated the experimental data. EE, IdlR, and MMA wrote the paper with inputs from all authors.

