## Supplementary Figures for "RATTLE: Reference-free reconstruction and quantification of transcriptomes from Nanopore sequencing"

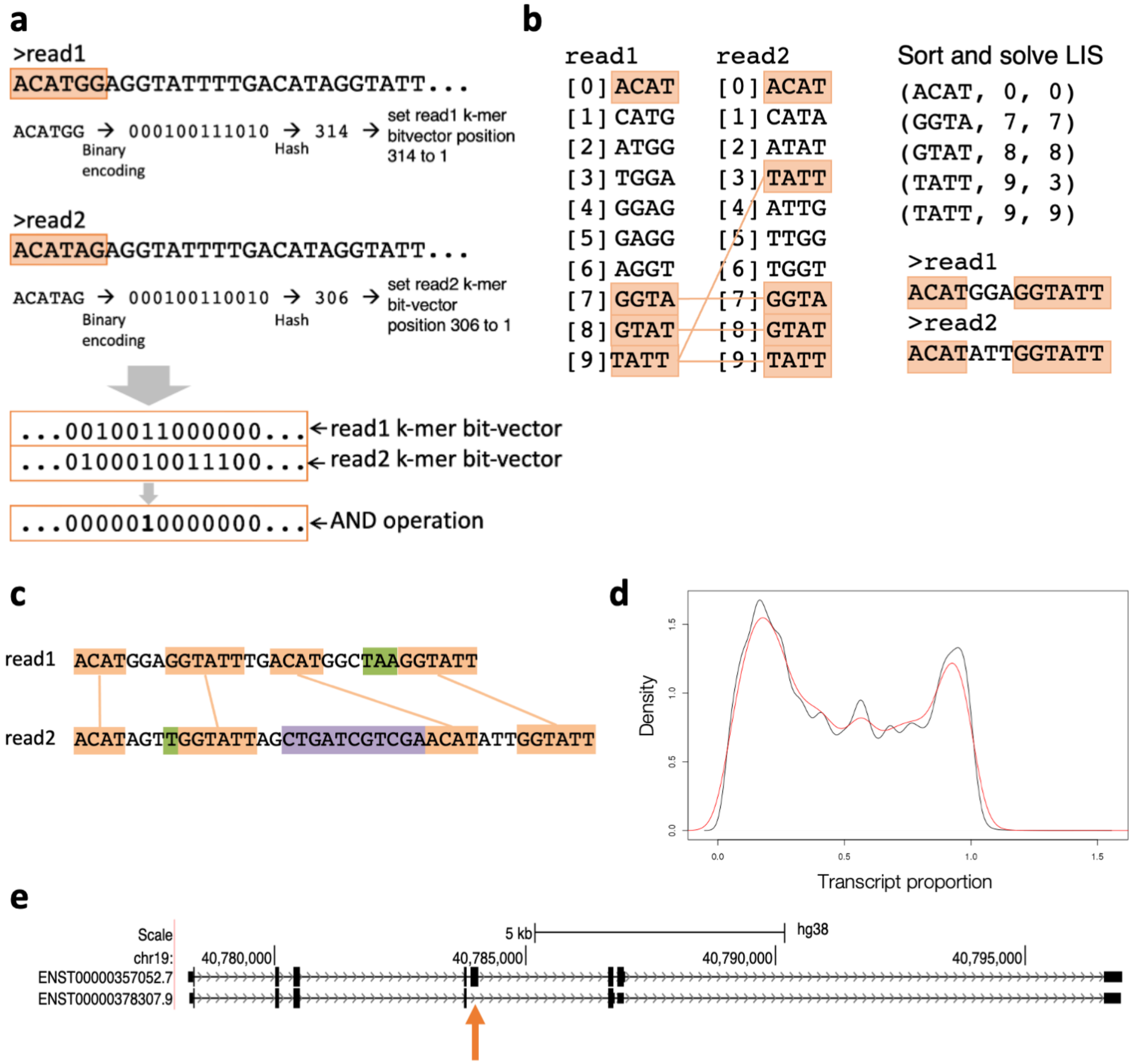


**Supplementary Figure 1.** RATTLE similarity between two reads is calculated in two steps. In the first step **(a),** RATTLE computes a binary representation of the k-mer content in each read (bit-vector) using hashing and calculating the number of common k-mers using an AND operation with the bit-vectors. The first similarity score is the fraction of unique k-mers in common over the maximum of unique k-mers in the two reads. If this score exceeds a predefined threshold, a second comparison is performed. In this second step **(b)**, all k-mers in both reads and their positions are extracted. This list is sorted by the positions in the first read, and the Longest Increasing Subsequence (LIS) problem is solved with a dynamic programming algorithm for the position of the k-mers in the second read. This yields a common set of co-linear k-mers between two reads. The second similarity value is defined as the number of bases covered by these k-mers over the length of the shortest read in the pair, and this defines the RATTLE score. **(c)** Illustration of how two reads in a gene cluster are separated into transcript clusters. Blocks of adjacent matching k-mers (in orange) are separated by gaps. Length differences in these gaps may be due to base-calling errors (in green) or to different exonic content (in purple). The variance of these gap differences between reads from two different transcripts is expected to be larger than the variance between reads from the same transcript. A threshold for this variance can be set as an input parameter. **(d)** Proportion of the annotated (Gencode v29)

transcript length (x axis) covered by real cDNA reads (black line) and by reads simulated with DeepSimulator (Li et al. 2018) (red line). The x axis goes over 1 because some of the cDNA reads were longer than the transcript they were mapped to. The cDNA dataset was from the Nanopore sequencing consortium https://github.com/nanopore-wgs-consortium/NA12878 - John Hopkins replicate 1. **(e)** We depict the exonic structure of the two transcripts, ENST00000378307.9 and ENST00000357052.7, used to compare the variance in the distribution of gap-length differences between adjacent matching k-mers (Fig. 1c). Reads were simulated from these two transcripts, which only differed by an alternative exon (highlighted with an orange arrow) of length 154nt.

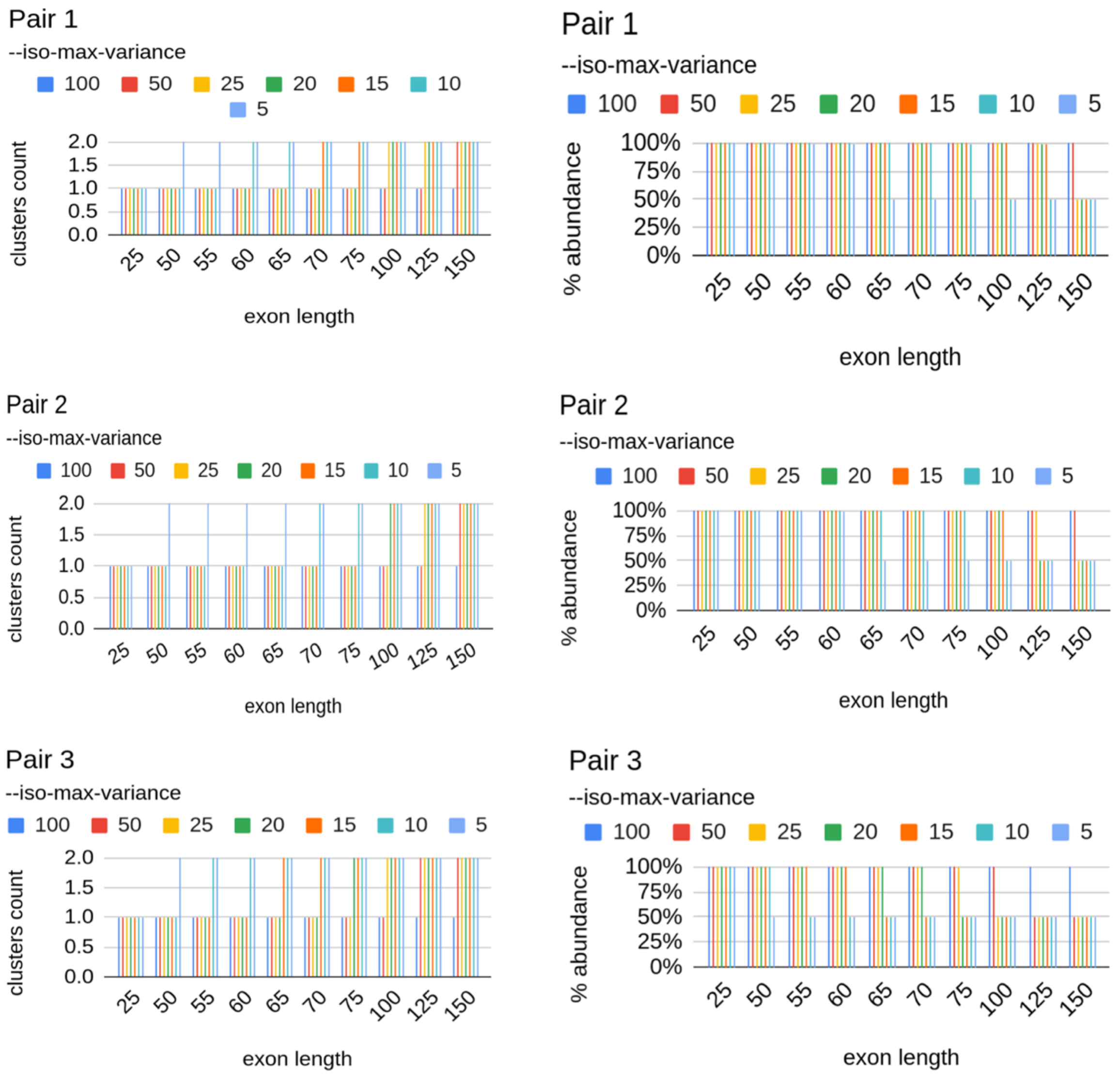


**Supplementary Figure 2.** We analysed three pairs of isoform transcripts: pair 1 (ENST00000378307.9, ENST00000357052.8), pair 2 (ENST00000667721.2, ENST00000684870.1), pair 3 (ENST00000525981.1, ENST00000526562.5). Each pair of isoforms differed from each other by one internal alternative exon (>150nt). We removed sequence from the alternative exon to obtain 10 variable exon lengths between 25 and 150: 25, 50, 55, 60, 65, 70, 75, 100, 125, 150. We used DeepSimulator to simulate 1000 Nanopore reads from each isoform. Then, we ran RATTLE and only changed --iso-max-variance parameter value from 5 to 100: 5, 10, 15, 20, 25, 50, 100. That is, we ran RATTLE for 10 x 7 = 70 different configurations for each isoform pair. We used the RATTLE default values for all other parameters (https://github.com/comprna/RATTLE). For all simulations, each corresponding to an alternative exon length and a –max-iso-variance value, RATTLE identified 1 single gene cluster. For each simulation, we further calculated the number of transcript clusters produced and the abundance of the most abundant transcript cluster. The abundance was calculated as the proportion of reads from the gene cluster that supports a given transcript. A perfect result would correspond to 2 transcript clusters and an abundance of 50% for the most abundant transcript (the other transcript having 50% too). The plotted data is available in Supp. Table S1.

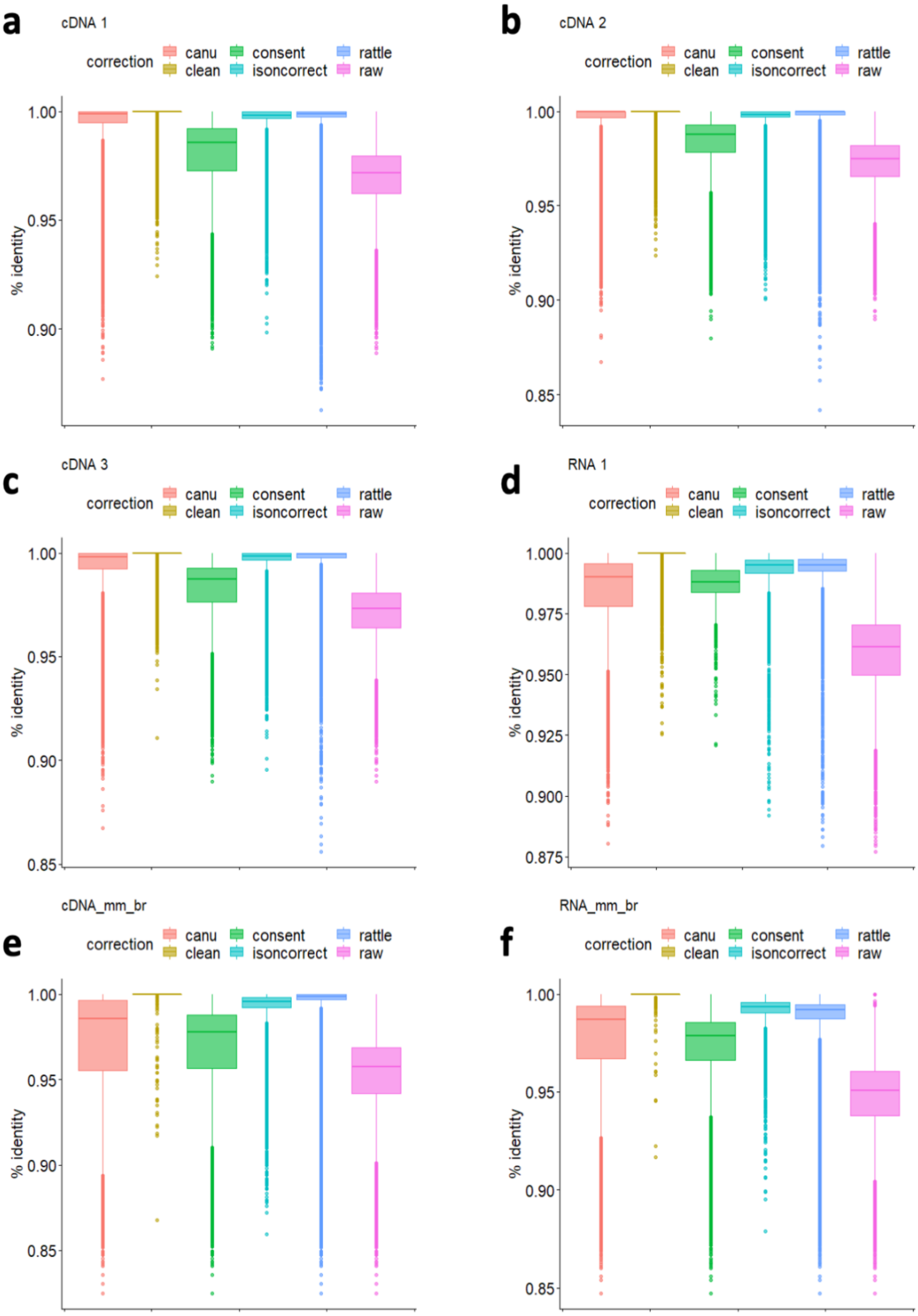


**Supplementary Figure 3.** Distribution of percentage identity for SIRV reads before (raw) and after correcting with RATTLE, CONSENT, Lorma, Canu and TranscriptClean (clean) for the samples of **(a)** cDNA1 (Human brain cDNA-seq), **(b)** cDNA2 (Human brain cDNA-seq), **(c)** cDNA3 (Human heart cDNA-seq), **(d)** RNA1 (Human heart direct RNA-seq), **(e)** cDNA_br_mm (Mouse brain cDNA-seq) (ERR2680377), **(f)** RNA_br_mm (Mouse brain RNA-seq) (ERR2680375). Percentage identity was calculated as the number of correct matches divided by the total length of the aligned region.

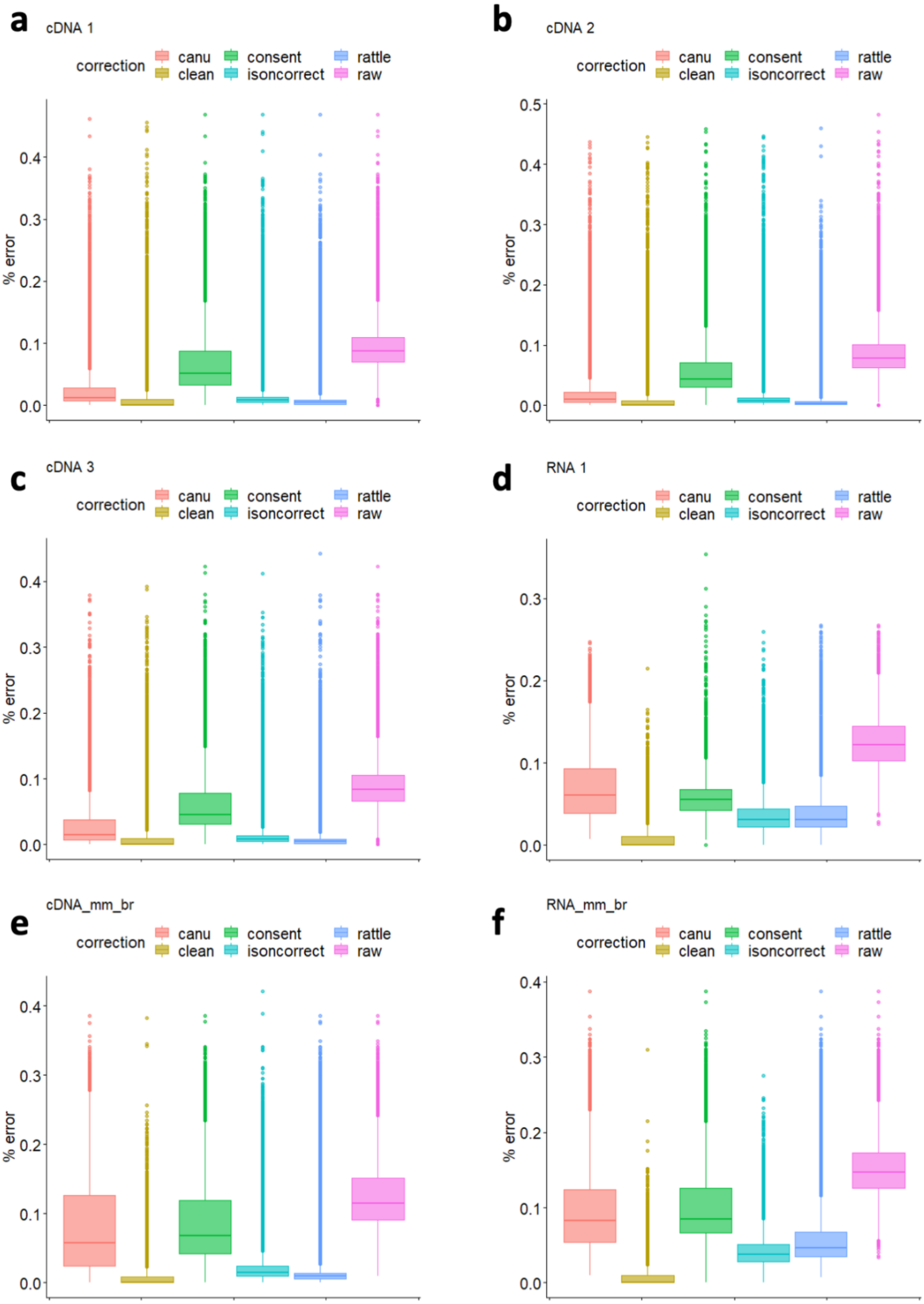


**Supplementary Figure 4.** Distributions of error rates for SIRV reads before (raw) and after correcting with RATTLE, CONSENT, Lorma, Canu and TranscriptClean (clean) for the same samples as Supp. Fig. 2: **(a)** cDNA1 (Human brain cDNA-seq), **(b)** cDNA2 (Human brain cDNA-seq), **(c)** cDNA3 (Human heart cDNA-seq), **(d)** RNA1 (Human heart direct RNA-seq), **(e)** cDNA_br_mm (Mouse brain cDNA-seq) (ERR2680377), **(f)** RNA_br_mm (Mouse brain RNA-seq) (ERR2680375). Error rate was calculated as the sum of insertions, deletions and substitutions divided by the length of the read.

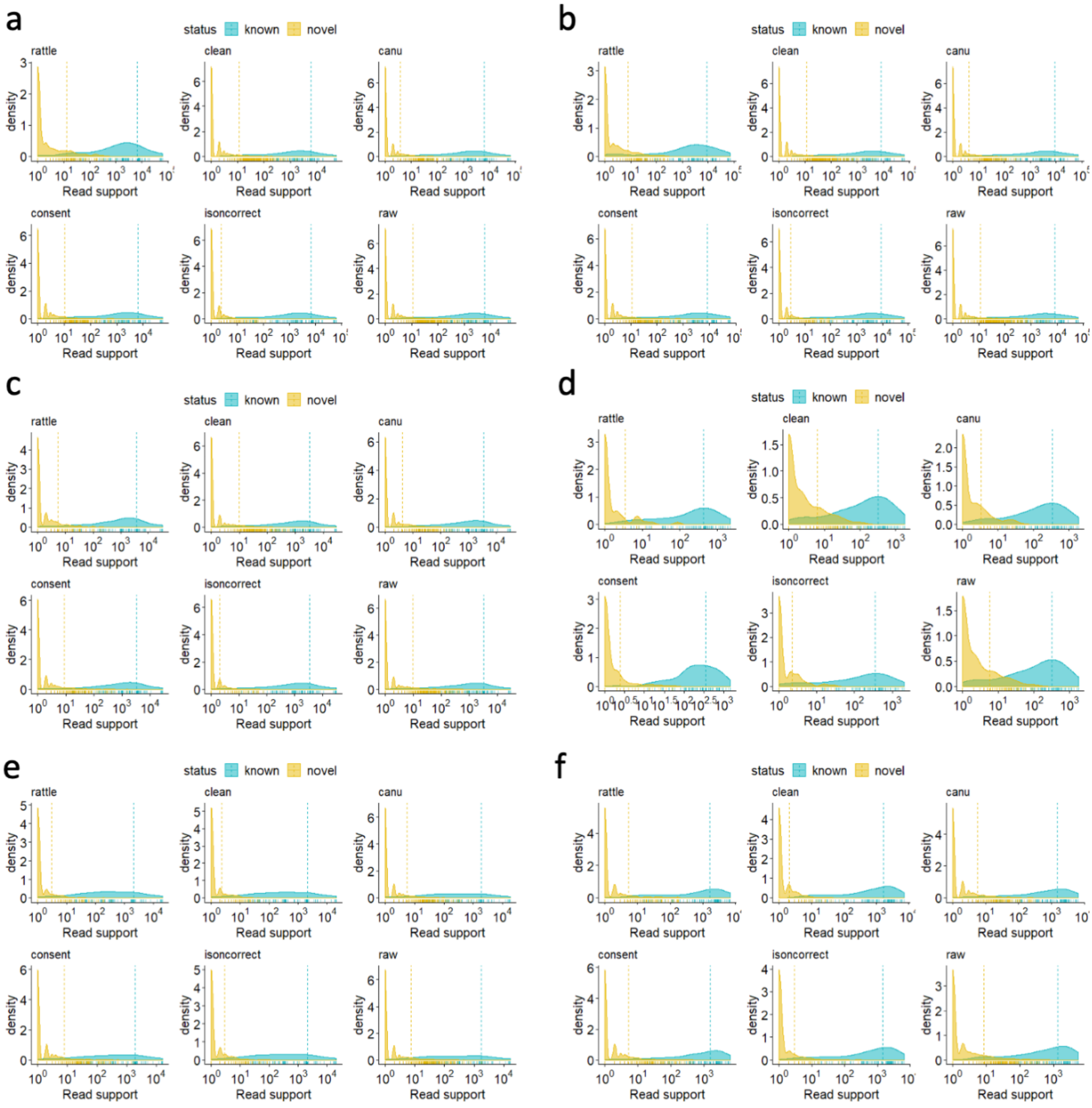


**Supplementary Figure 5.** Density profiles (y axis) for the read support (x axis) for true positives (known) and false positive (novel) introns using the SIRV annotations as reference, for RATTLE, CONSENT, Canu, isONcorrect, and TranscriptClean (clean) for all samples tested: **(a)** cDNA1 (human brain), **(b)** cDNA2 (human brain), **(c)** cDNA3 (human heart), **(d)** RNA1 (human heart), **(e)** Mouse brain Nanopore cDNA-seq (ERR2680377) (cDNA_br_mm), and **(f)** Mouse brain Nanopore RNA-seq (ERR2680375) (RNA_mm_br).

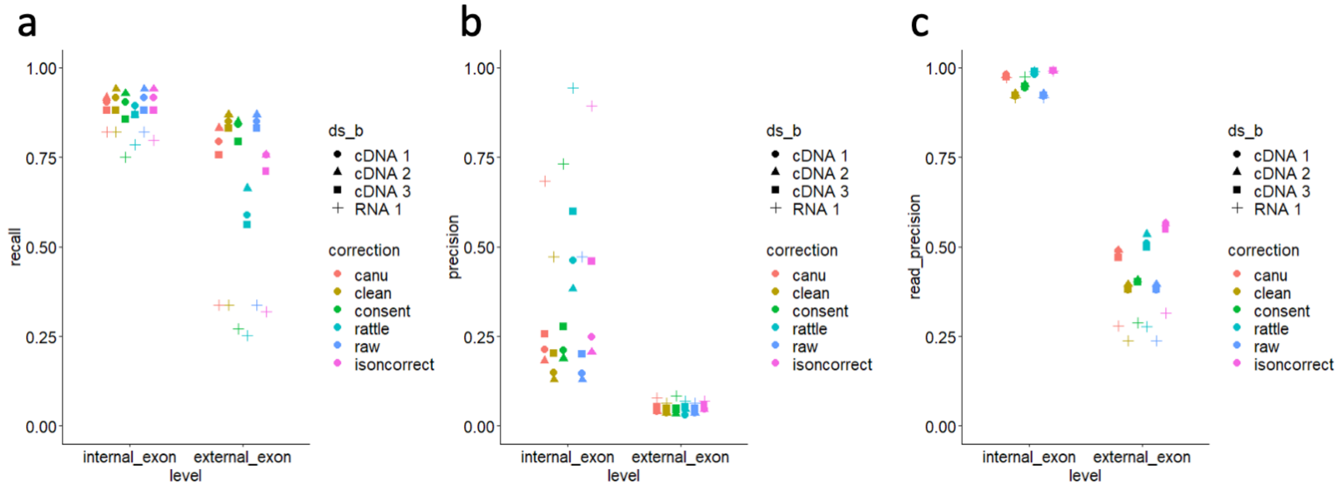


**Supplementary Figure 6. (a)** Recall of unique (internal and external) exons in SIRV transcript isoforms obtained by mapping reads to the SIRV genome before (raw) and after correction with RATTLE, CONSENT, Canu, isONcorrect, and TranscriptClean (clean) for the SIRV reads from the experiments run with the direct RNA protocol (RNA 1) and the (PCR-based) cDNA protocol (cDNA 1, 2, 3) (Methods). Recall was calculated as the fraction of unique annotated exons (internal or external) found by each method with 5 or more supporting reads. Exons were defined as external if they were first or last in any of the transcript isoforms; and internal, otherwise. **(b)** Precision of unique (internal and external) exons for the same methods and datasets as (a). Precision was calculated as the fraction of unique exons predicted by reads that matched correctly the annotation and had support of 5 or more reads. **(c)** Read-precision for the (internal and external) exons for the same methods and datasets as in (a). Read-precision was calculated as the fraction from the total number of exons predicted in reads that corresponded to annotated exons, and counting only those predicted by 5 or more reads.

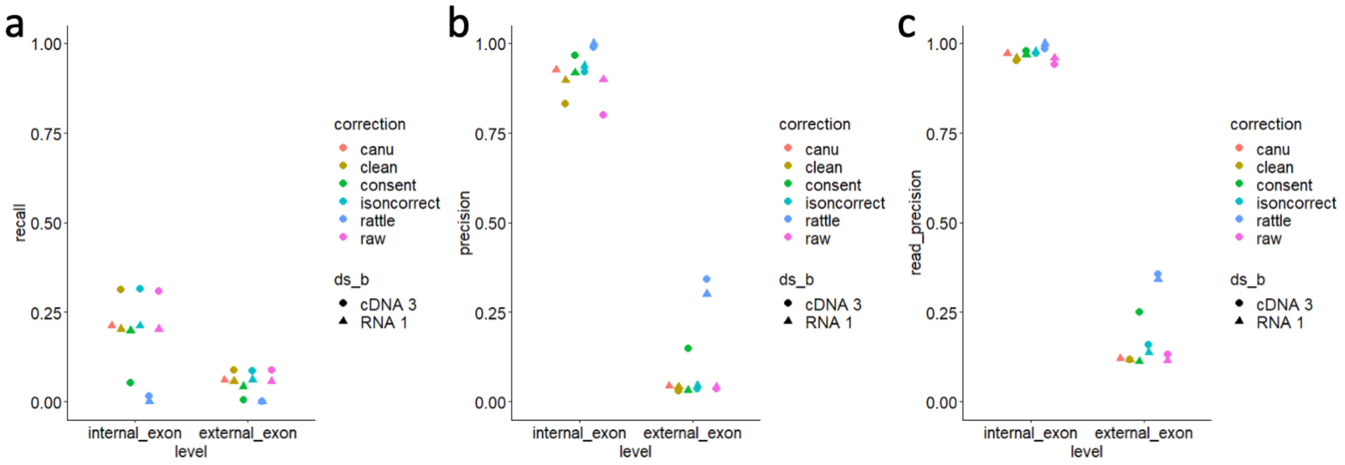


**Supplementary Figure 7. (a)** Recall of unique (internal and external) exons in expressed Gencode transcript isoforms (>5 reads) obtained by mapping reads to the genome before (raw) and after correction with RATTLE, CONSENT, Canu, isONcorrect and TranscriptClean (clean) for all (non-SIRV) reads from the experiments for heart direct RNA protocol (RNA1) and the (PCR-based) cDNA protocol (cDNA3). We did not include the other datasets as some of the other methods were not able to conclude the analysis. Recall was calculated as the fraction of unique annotated exons (internal or external) found by each method with 5 or more supporting reads. Exons were defined as external if they were first or last in any of the transcript isoforms; and internal, otherwise. **(b)** Precision of unique (internal and external) exons for the same methods and datasets as (a). Precision was calculated as the fraction of unique exons predicted by reads that correctly matched the annotation and had support of 5 or more reads. **(c)** Read precision for the (internal and external) exons for the same methods and datasets as in (a). Read precision was calculated as the fraction from the total number of exons predicted in reads that corresponded to annotated exons, and counting only those predicted by 5 or more reads.

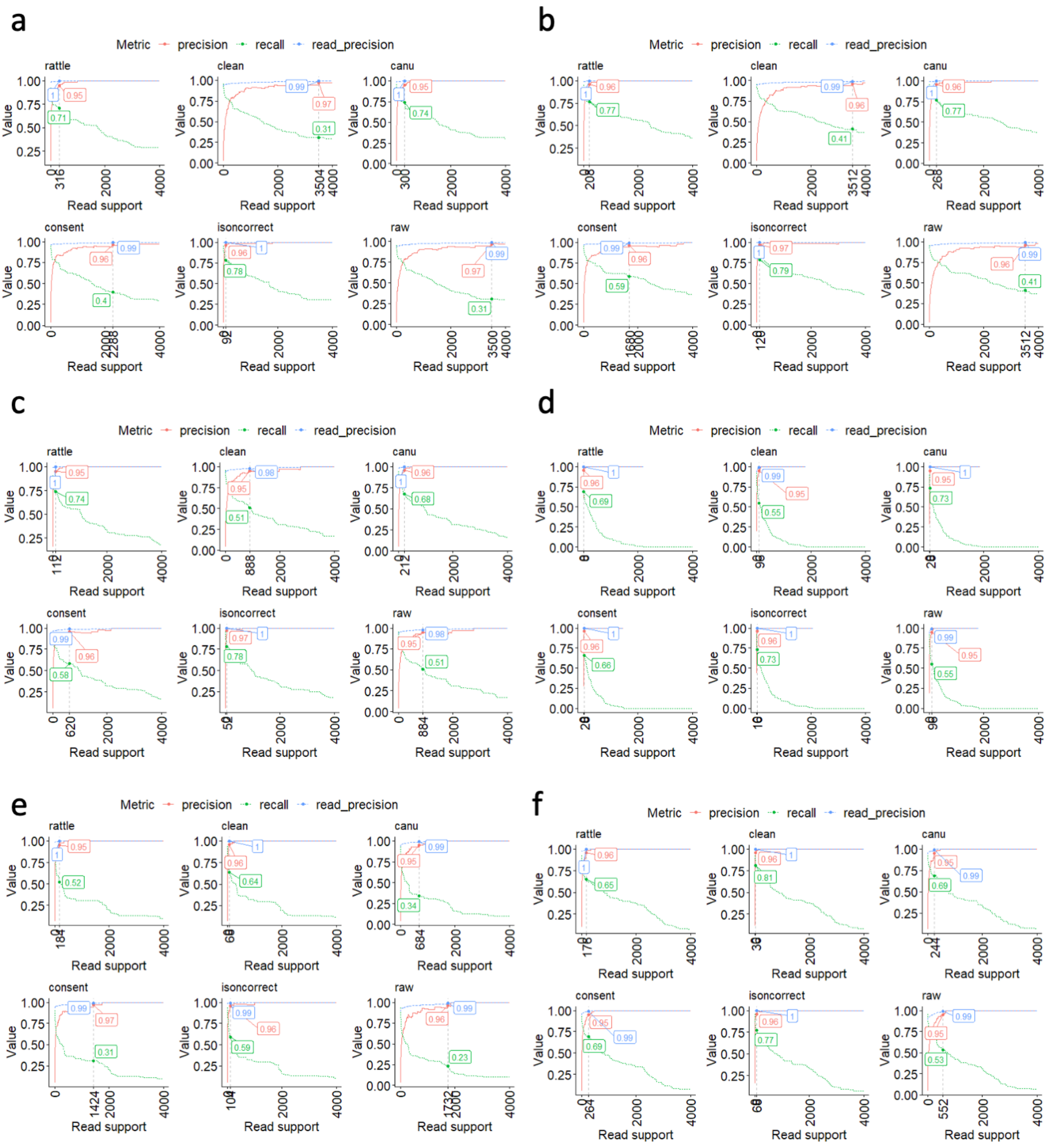


**Supplementary Figure 8.** We plot the recall (green), precision (red) and read precision (blue) of the SIRV introns as a function of an expression cut-off (x axis) given in terms of the number of reads supporting the introns. We indicate for each case the threshold at which a precision of approximately 0.95 is achieved. For that threshold we indicate the corresponding recall, precision, and read-precision values. The plot corresponds to the samples **(a)** cDNA1 (Human brain cDNA-seq), **(b)** cDNA2 (Human brain cDNA-seq), **(c)** cDNA3 (Human heart cDNA-seq), **(d)** RNA1 (Human heart direct RNA-seq), **(e)** Mouse brain Nanopore cDNA-seq (ERR2680377) (cDNA_br_mm), **(f)** Mouse brain Nanopore RNA-seq (ERR2680375) (RNA_br_mm).

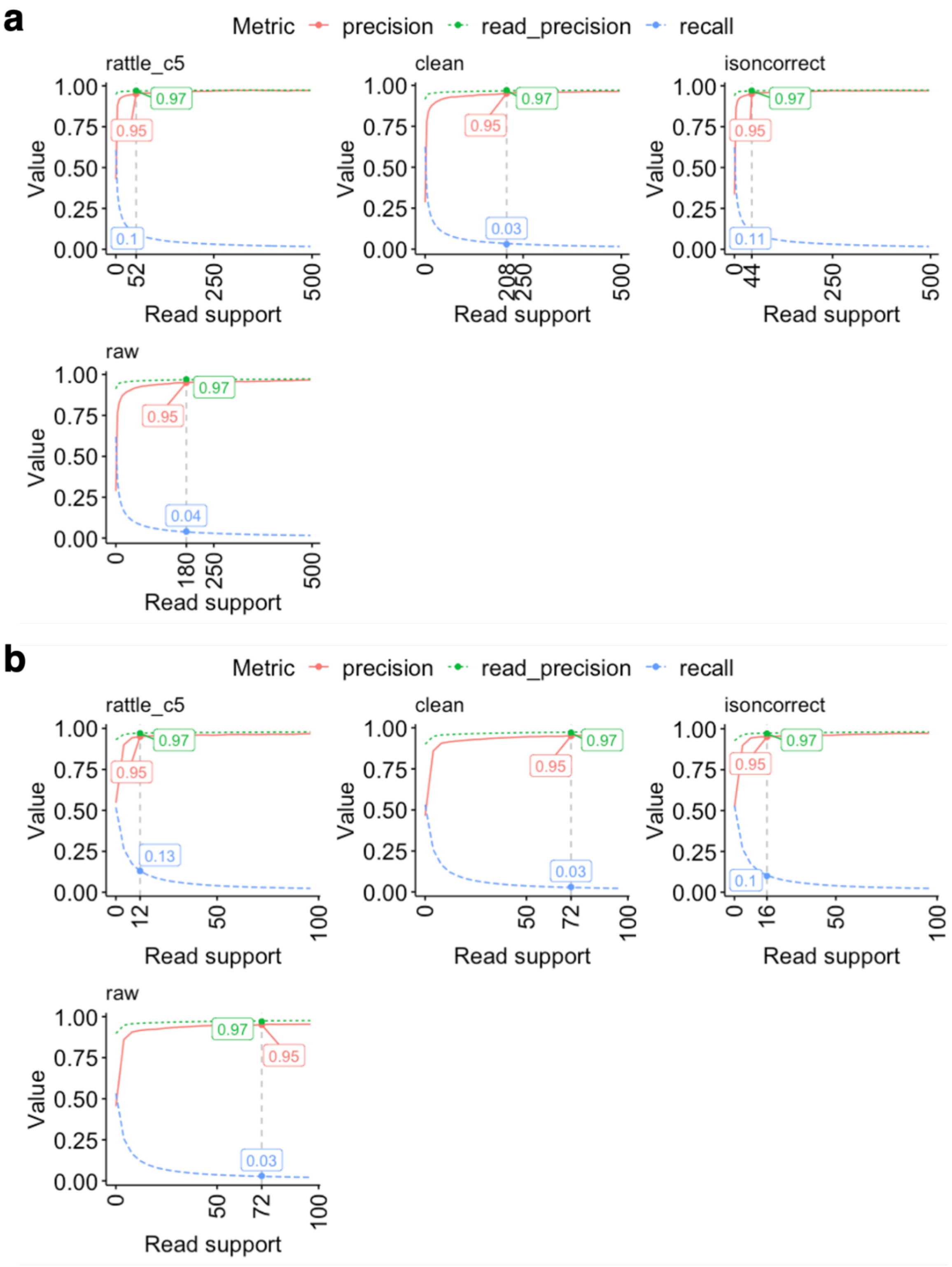


**Supplementary Figure 9.** We plot the recall (green), precision (red) and read-precision (blue) of the Gencode introns as a function of an expression cut-off (x axis) given in terms of the number of reads supporting the introns. We used introns from transcripts with expression evidence (>5 reads). We indicate for each case the read support at which a precision of approximately 0.95 is achieved. For that threshold we indicate the corresponding recall, precision, and read-precision values. The plot corresponds to the samples Human heart cDNA-seq (cDNA3) **(a)** and dRNA-seq (RNA1) **(b)**. For RATTLE, we used the parameter configuration c5 (see Supplementary Table S10).

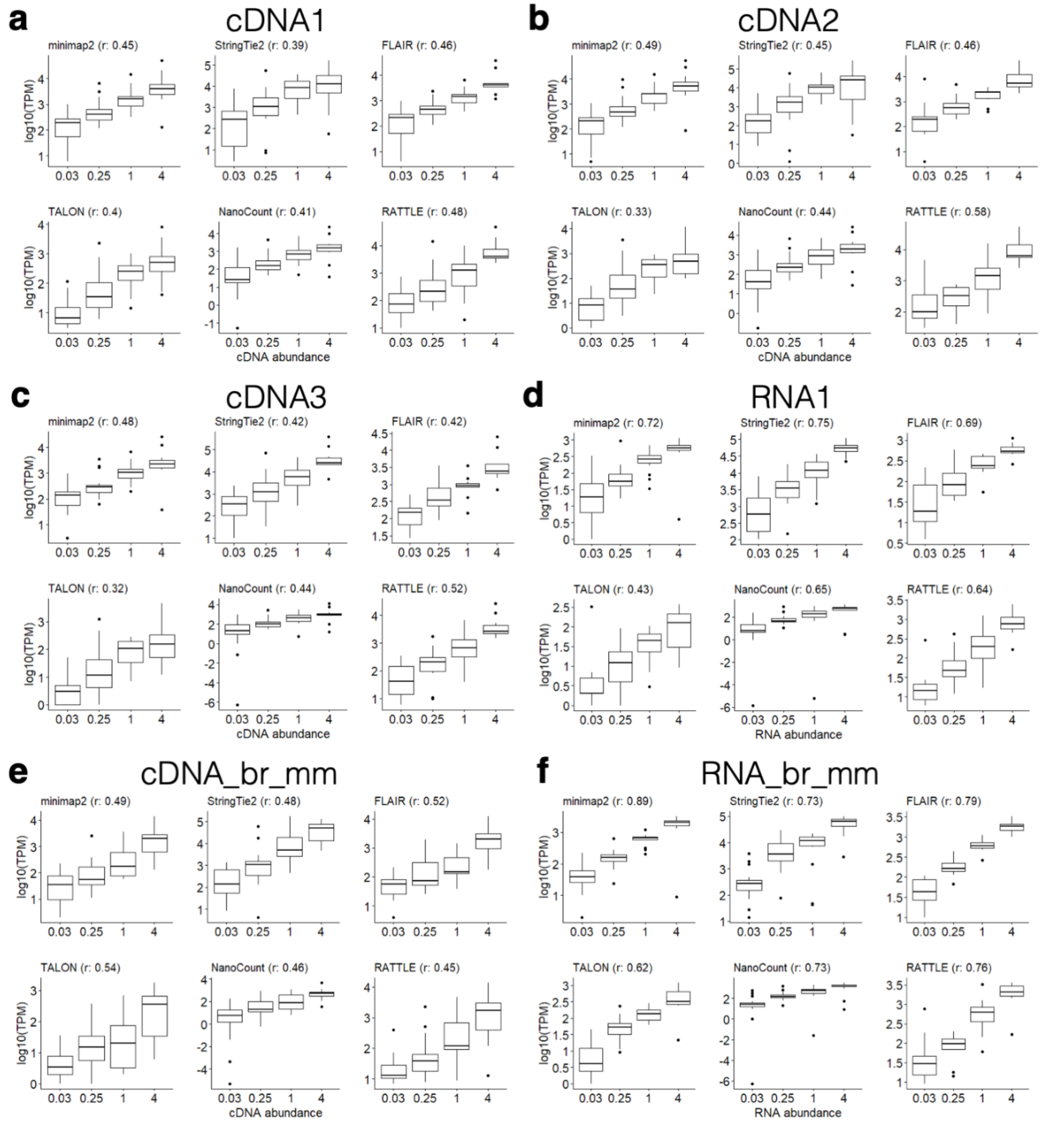


**Supplementary Figure 10.** Comparison of predicted (y axis) and known (x axis) abundances for the SIRV transcript isoforms. We show the predictions by RATTLE, FLAIR, StringTie2, TALON. We also considered abundances estimated by counting the best read mappings with Minimap2 (minimap2), and the abundance estimated with NanoCount (https://github.com/a-slide/NanoCount), which uses an expectation maximization (EM) algorithm. We show the Pearson correlation *r* for each method. Units on the y-axis vary according to method: RATTLE provides abundances in terms of read counts per million, similar to TALON and FLAIR. StringTie2 produces a TPM value using the same formula as for short reads. For NanoCount and minimap2 we give read counts per million. We show the results for cDNA and dRNA samples: **(a)** cDNA1 (human brain), **(b)** cDNA2 (human brain), **(c)** cDNA3 (human heart), **(d)** RNA1 (human heart), **(e)** cDNA_br_mm (mouse brain), and (f) RNA_br_mm (mouse brain).

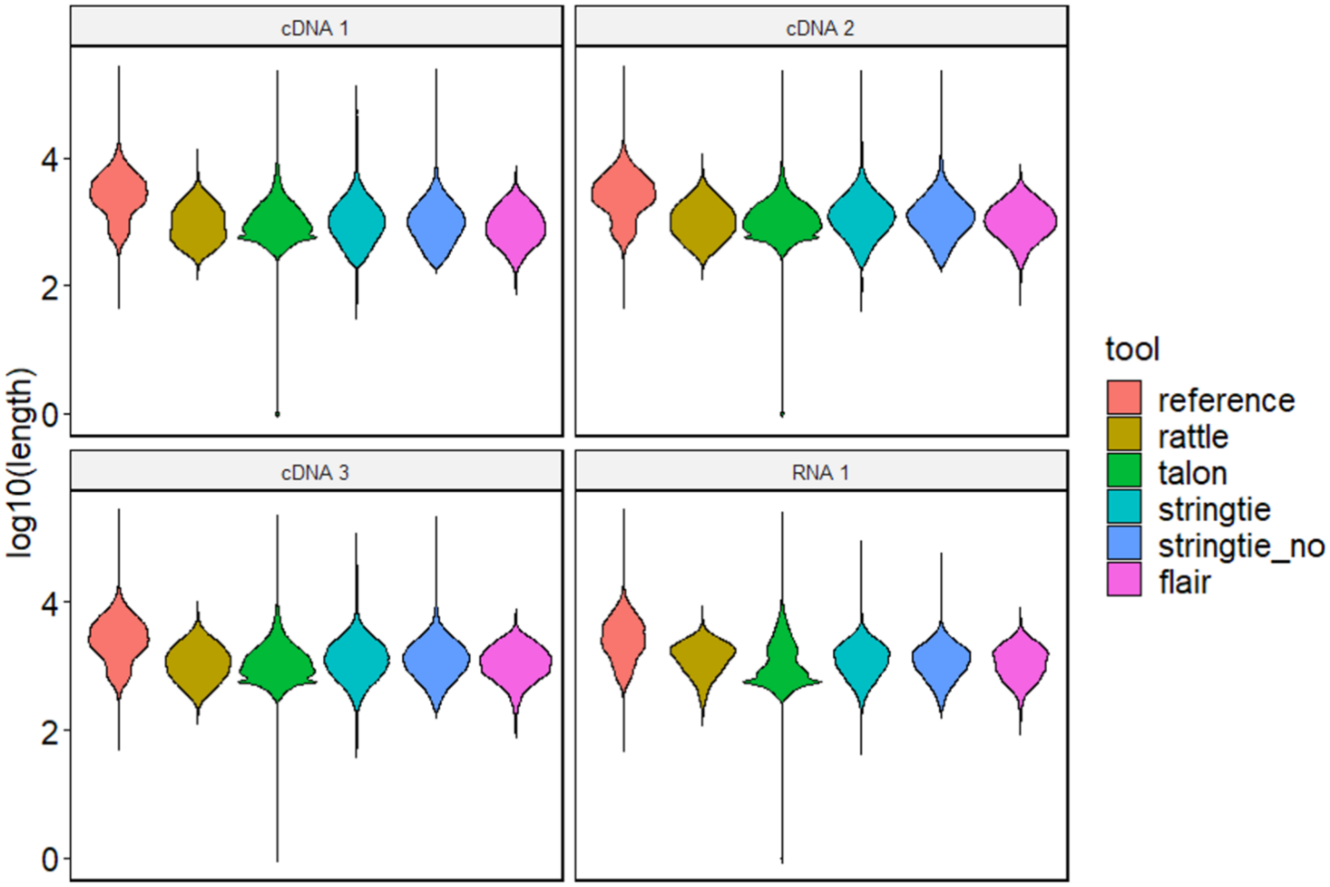


**Supplementary Figure 11.** Distribution of the lengths (y-axis, log10 scale) of the transcripts predicted by each method compared with the transcripts from the reference (Gencode v29) in the three cDNA and one RNA samples analysed. StringTie2 was run with (stringtie) and without (stringtie_no) using the annotation. We considered only predicted and annotated transcripts with support of more than 5 reads.

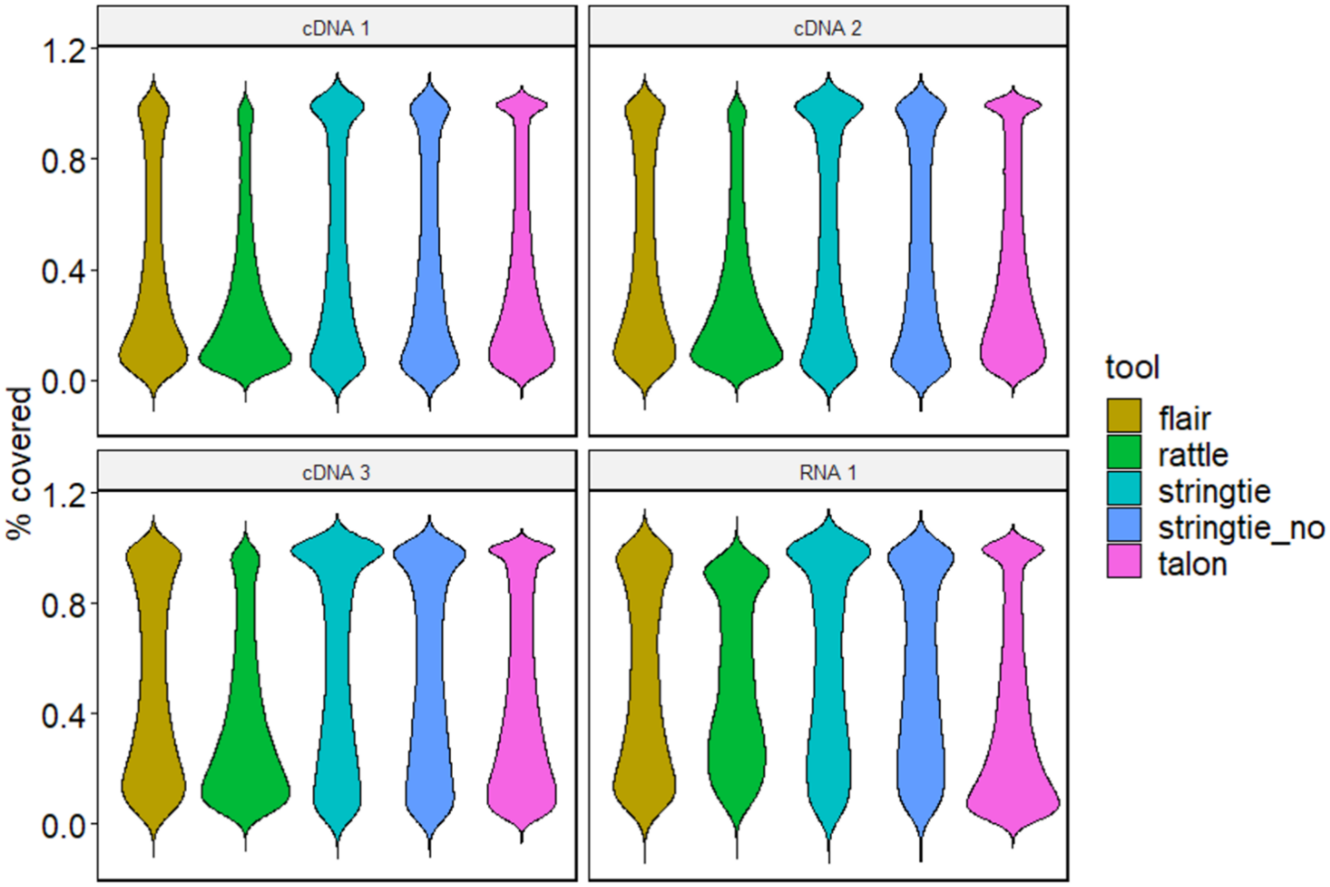


**Supplementary Figure 12.** Distributions of the percentage of the reference transcript covered (y-axis) by the predicted transcript with the best match to it, for each of the methods tested. For StringTie2 we include the version with (stringtie) and without the annotation (stringtie_no).

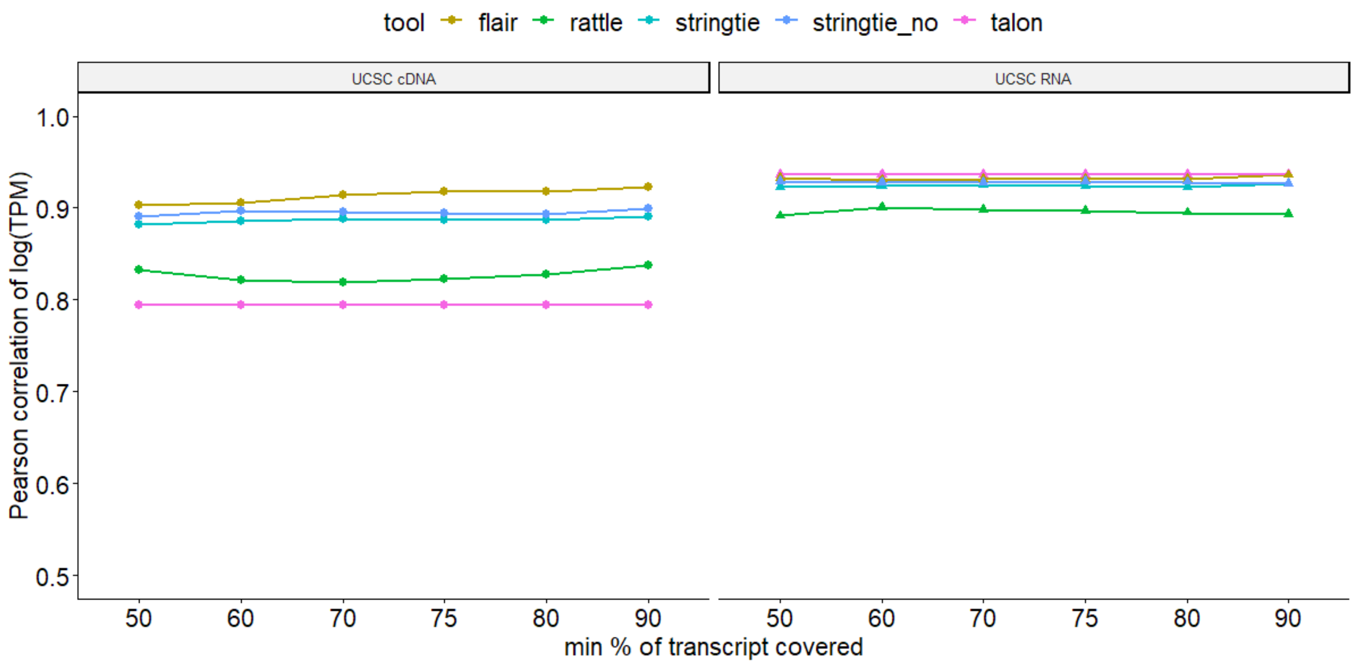


**Supplementary Figure 13.** For each minimum coverage threshold (x axis), we plot the Pearson correlation (y axis) of the abundances of the predicted transcripts and the assigned annotated transcripts based on such minimum coverage threshold. We give this correlation for cDNA data (left panel) and dRNA data (right panel). A minimum coverage of m% means that the best mapping for each predicted transcript covers an annotated transcript at least for m% of its length. Coverage is calculated as the number of matches plus substitutions divided by the target length, in a percentage scale.

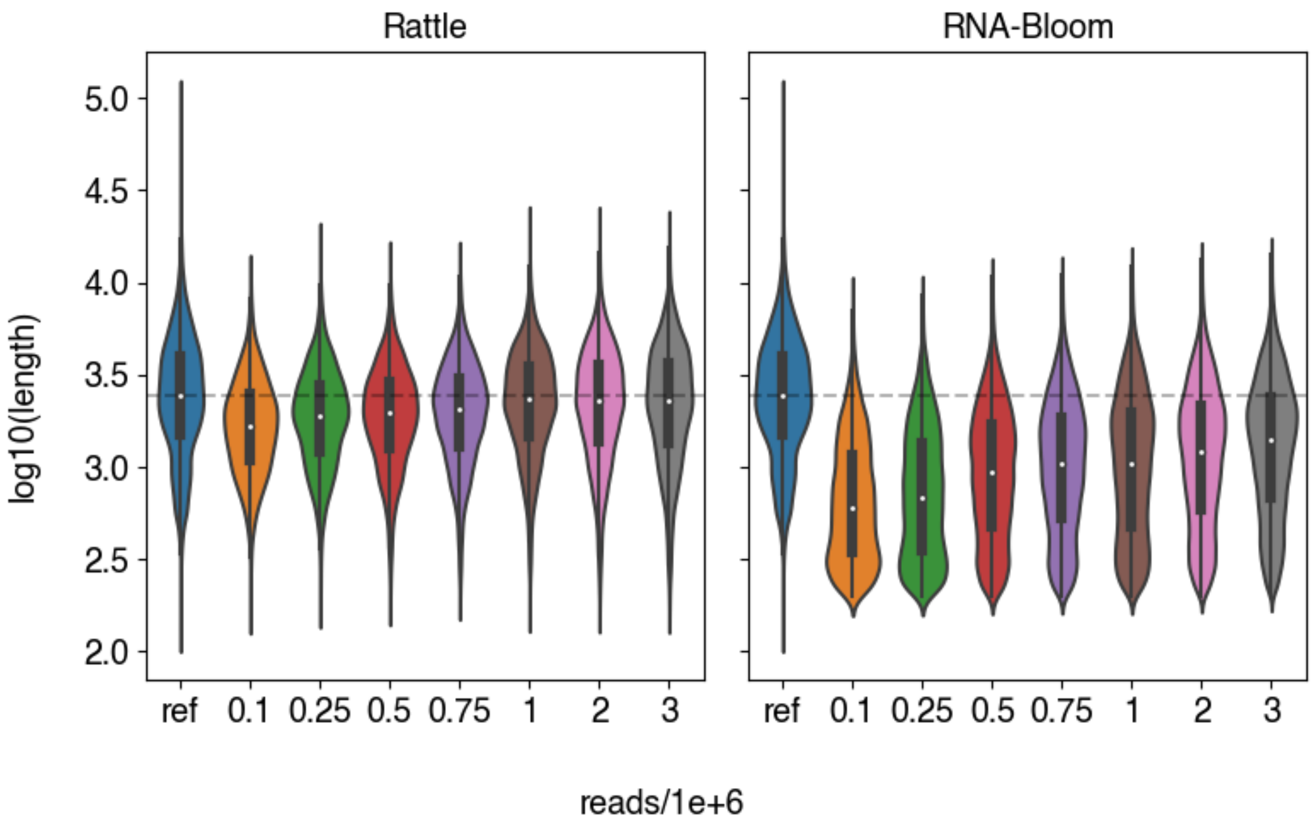


**Supplementary Figure 14.** Distribution of the lengths (y-axis, log10 scale) of the transcripts predicted by RATTLE (left) and RNA-Bloom (right) for an increasing number of input reads (x axis): 100k, 250k, 500k, 750k, 1M, 2M, and 3M reads. The distributions are compared with the length distribution of the annotation (ref), consisting of all Ensembl annotated transcripts (after discarding pseudogenes and short non-coding RNAs) with evidence of expression using Minimap2 (options: -t 24 -x map-ont --secondary=no) . Nanopore data was direct RNA sequencing from HEK293 wild type cells from the European Nucleotide Archive (ENA), accession PRJEB40872.

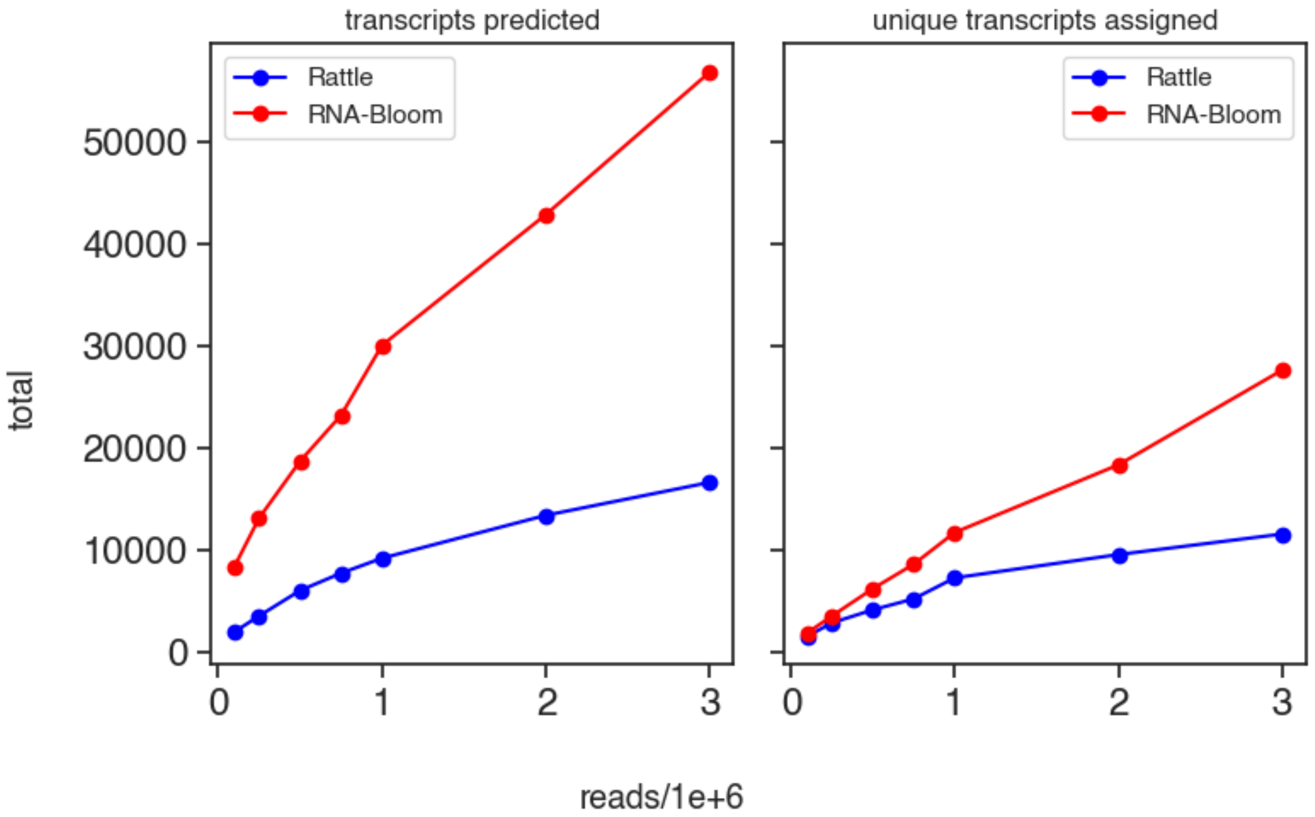


**Supplementary Figure 15.** Saturation plot of transcripts using an incremental number of dRNA reads used as input: 100k, 250k, 500k, 750k, 1M, 2M, and 3M reads. Left panel: for each input (x-axis), we give the total number of transcripts predicted by each method (y-axis). Right panel: for each input (x-axis), we give the total number of annotated transcripts matched by the predicted transcripts from the left panel with a coverage of at least 75%. Nanopore data was direct RNA sequencing from HEK293 wild type cells from the European Nucleotide Archive (ENA), accession PRJEB40872.
